# Dynamic sensitivity analysis of a mathematical model describing the effect of the macroalgae *Asparagopsis taxiformis* on rumen fermentation and methane production under *in vitro* continuous conditions

**DOI:** 10.1101/2024.06.19.599712

**Authors:** Paul Blondiaux, Tristan Senga Kiessé, Maguy Eugène, Rafael Muñoz-Tamayo

## Abstract

Ruminants play an important role in global warming by emitting enteric methane (CH_4_) through the degradation of feeds by the rumen microbiota. To better understand the dynamics fermentation outputs, including CH_4_ and volatile fatty acids (VFA) production, mathematical models have been developed. Sensitivity analysis (SA) methods quantify the contribution of model input parameters (IP) to the variation of an output variable of interest. In animal science, SA are usually conducted in static condition. In this work, we hypothesized that including the dynamic aspect of the rumen fermentation to SA can be useful to inform on optimal experimental conditions aimed at quantifying the key mechanisms driving CH_4_ and VFA production. Accordingly, the objective of this work was to conduct a dynamic SA of a rumen fermentation model under *in vitro* continuous conditions (close to the real *in vivo* conditions). Our model case study integrates the effect of the macroalgae *Asparagopsis taxiformis* (AT) on the fermentation. AT has been identified as a potent CH_4_ inhibitor via the presence of bromoform, an anti-methanogenic compound. We computed Shapley effects over time for quantifying the contribution of 16 IPs to CH_4_ (mol/h) and VFA (mol/l) variation. Shapley effects integrate the three contribution types of an IP to output variable variation (individual, via the interactions and via the dependence/correlation). We studied three diet scenarios accounting for several doses of AT relative to Dry Matter (DM): control (0% DM of AT), low treatment (LT: 0.25% DM of AT) and high treatment (HT: 0.50% DM of AT). Shapley effects revealed that hydrogen (H_2_) utilizers microbial group via its Monod H_2_ affinity constant highly contributed (> 50%) to CH_4_ variation with a constant dynamic over time for control and LT. A shift on the impact of microbial pathways driving CH_4_ variation was revealed for HT. IPs associated with the kinetic of bromoform utilization and with the factor modeling the direct effect of bromoform on methanogenesis were identified as influential on CH_4_ variation in the middle of fermentation. Whereas, VFA variation for the three diet scenarios was mainly explained by the kinetic of fibers degradation, showing a high constant contribution (> 30%) over time. The simulations computed for the SA were also used to analyze prediction uncertainty. It was related to the dynamic of dry matter intake (DMI, g/h), increasing during the high intake activity periods and decreasing when the intake activity was low. Moreover, CH_4_ (mol/h) simulations showed a larger variability than VFA simulations, suggesting that the reduction of the uncertainty of IPs describing the activity of the H_2_ utilizers microbial group is a promising lead to reduce the overall model uncertainty. Our results highlighted the dynamic nature of the influence of metabolic pathways on CH_4_ productions under an anti-methanogenic treatment. SA tools can be further exploited to design optimal experiments studying rumen fermentation and CH_4_ mitigation strategies. These optimal experiments would be useful to build robust models that can guide the development of sustainable nutrition strategies.

## 1. Introduction

Reducing methane (CH_4_) emissions from ruminants is an important challenge for the livestock sector. At the global level, these emissions are responsible of 14.5% of total greenhouse gases (GHG) from human activity sources (FAO, 2017), highlighting the important role of ruminants in global warming. In this context, Masson-Delmotte et al. (2019) highlighted that decreasing agricultural CH_4_ emissions by 11 to 30% of the 2010 level by 2030 and by 24 to 47% by 2050 must be achieved to meet the 1.5 °C target of the Paris Agreement. In addition, Arndt et al. (2022) indicated that some mitigation strategies scenarios may allow to meet the 1.5 °C target by 2030. However, this study also highlighted that it was not possible to meet this target by 2050 considering that expected increase in milk and meat demand will lead to an increase of GHG emissions.

Ruminants produce CH_4_ during the degradation and fermentation of feeds (Morgavi et al., 2010; Beauchemin et al., 2020). The fermentation process is done by a complex community of microbes inhabiting the forestomach (rumen) of ruminants. These microbial community (rumen microbiota) are constituted by members of bacteria, archaea, fungi and protozoa. The products of fermentation include volatile fatty acids (VFA), which are useful compounds for the animal, and CH_4_. The development of mitigation actions aiming to reduce the enteric CH_4_ production without affecting animal performance and welfare is a crucial challenge for the field (Hristov et al., 2013; Pellerin et al., 2013; Torres et al., 2020).

To better understand rumen fermentation and help design such strategies, mechanistic models describing the dynamic process of the rumen fermentation were developed. A synthesis of the characteristics of these models is presented by Tedeschi et al. (2014). Among these, the three most popular dynamic mechanistic models of rumen fermentation are: Molly (Baldwin et al., 1987), Dijkstra et al. (1992) and Karoline (Danfær et al., 2006). More recently, Pressman and Kebreab (2024) provided a review of current state of rumen models, and tried to identify future needs for improving the representation of the impact of feed additives on rumen fermentation dynamic. This work added the model of Muñoz-Tamayo et al. (2021, 2016) as another “lineage” of rumen models. It is one of the two mechanistic models available representing the effect of a feed additive on the rumen fermentation.

All the mechanistic models involved numerous input parameters (IPs) representing biological and physical processes. The complexity of such models raises the need to investigate model behaviour, including the various relationships among IPs and outputs. To address this need, sensitivity analysis (SA) methods were used to assess the contribution of IP variability on the variability of the output of interest, identifying IPs which contribute the most to model predictions variability from those having a negligible effect (Faivre et al., 2013; Iooss and Lemaître, 2015; Saltelli et al., 2008, 2005).

In animal nutrition, SA is usually conducted on mechanistic models with the main objective of reducing their complexity and identifying which IPs requires more accurate measurements for reducing output uncertainty. For instance, Huhtanen et al. (2015) and van Lingen et al. (2019) used linear regressions for describing the effects of some parameters on the variation of daily scale enteric CH_4_ emissions in the Karoline model and an updated version of the Dijkstra model, respectively. In addition, Morales et al. (2021) and Dougherty et al. (2017) computed the Sobol indices (Sobol, 1993) for quantifying the effects of 19 and 20 parameters on several output variables of the Molly and AusBeef (Nagorcka et al., 2000) models, respectively. Morales et al. (2021) did not consider the CH_4_ production among the 27 output variables studied, while VFAs were considered. Dougherty et al. (2017) considered the daily CH_4_ production in the output variables. In both studies, uniform distributions were set for exploring parameter variability. Recently, Merk et al. (2023) adapted and calibrated the model of Muñoz-Tamayo et al. (2021) to represent experimental data from the *in vitro* RUSITEC study of Roque et al. (2019), which aimed at evaluating the effect of the macroalgae *Asparagosis taxiformis* (AT) on CH_4_ production and rumen microbiota. Authors used a Sobol based approach for identifying key parameters associated to microbial pathways driving CH_4_ production with and without the presence of AT. Despite these contributions, SA is rarely applied to animal nutrition models, but many applications are found in other field of animal science. For instance, many works implementing SA on epidemiological models are available in the literature. Lurette et al. (2009) conducted a dynamic SA based on a principle component analysis and on analysis of variance, to identify key parameters influencing *Salmonella* infection dynamics in pigs. More recently, Farman et al. (2025) investigated the impact of various parameters on the progression of the brucellosis infection disease in cattle, by computing the partial rank correlation coefficient.

Although different SA approaches were applied to mechanistic models of the rumen fermentation in the literature, several aspects still need to be explored. One aspect is the dynamic characteristic of the rumen fermentation. Most of the studies mentioned above explored sensitivity of rumen mechanistic models at a single time point or under steady state conditions. However, our hypothesis is that the influence of model parameters is of dynamic nature, since the rumen is a dynamic system. Characterizing such dynamic influence is of relevance to better understand rumen function (Morgavi et al., 2023), and represents the central step of an optimal experiment design aiming at identifying experimental conditions enabling accurate estimation of model parameters, and thus accurate characterization of mechanisms. Muñoz-Tamayo et al. (2014) highlighted the importance of optimal experimental designs for improving parameter estimation accuracy of a microalgae growth model. This characterization could also be useful to help design CH_4_ mitigation strategies, in a context where investigating the combined effect of several CH_4_ mitigating compounds is an active research topic in animal nutrition. To our knowledge, SA has not been applied in dynamic conditions for studying CH_4_ and VFA predictions of rumen models, highlighting a first gap to be filled.

Other aspect to explore in mechanistic models is the nature of the contribution of the IPs to model outputs by identifying: 1) the effect due to the IPs alone, 2) the effect due to the interactions between the IPs and 3) the effect due to the dependence or correlation between the IPs. Some references mentioned above implemented a method differentiating some effects of the contribution of an IP to output (individual, interaction and dependence/correlation). Dougherty et al. (2017) computed first-order and total Sobol indices (Homma and Saltelli, 1996), quantifying the individual and interaction effects of IPs. Whereas, van Lingen et al. (2019) concluded that there was no interaction between parameter covariates when studying the variation of daily scale enteric CH_4_ emissions. The quantification of the contribution due to the dependence/correlation between the IPs has been an important research activity of the applied mathematics field for several years now (Kucherenko et al., 2012; Mara et al., 2015; Xu and Zdzislaw Gertner, 2008). Not all the SA methods are able to identify these three effects, conducting to biases in the estimated sensitivity indices. The characterization of these three effects in rumen dynamic models was not performed in previous works, highlighting a second gap to be filled.

Therefore, the aim of this work was to fill these two gaps, by conducting a complete dynamic SA of a mechanistic model of rumen fermentation under *in vitro* continuous conditions accounting for the effect of AT on the fermentation and CH_4_ production. The representation under *in vitro* conditions means that the rumen fermentation is reproduced outside the living organism, in a controlled laboratory reactor.

The model studied extends previous developments of Muñoz-Tamayo et al. (2016, 2021). The AT macroalgae has been identified as a potent CH_4_ inhibitor (Machado et al., 2014), with reported *in vivo* reductions of CH_4_ emissions over 80% and 98% in beef cattle (Kinley et al., 2020; Roque et al., 2021). Moreover, developing dynamic models able to represent the effect of feed additives on rumen fermentation was highlighted as crucial by Pressman and Kebreab (2024), and AT is one of the most promising inhibitors. The original model (Muñoz-Tamayo et al., 2021) represented the fermentation under batch conditions. We extended the model to account for continuous conditions which aimed at providing a model closer to the *in vivo* conditions, which means a representation of the rumen fermentation integrating the animal. This extension allowed to simulate dietary scenarios accounting for several doses of AT. These scenarios were set according to an *in vitro* study (Chagas et al., 2019) aimed to reproduce forage-based diets typical of dairy and beef cattle systems, testing several doses of AT. We hypothesized that applying dynamic SA to these scenarios could be a way to improve our knowledge of microbial pathway mechanisms according to different diets and treatments.

The SA method implemented was the Shapley effects (Owen, 2014). The added value of the Shapley effects is the consideration of the three effects of an input on an output. Here, this is an academic exercise, as no dependencies or correlations between parameters are present in the model studied, but using this type of methods is crucial in the future. By applying this SA method over time in a context considering the effect of a CH_4_ inhibitor, we are able to identify the parameters explaining the variation of CH_4_ and VFA, and to investigate how the contribution of these parameters move according to the inhibitor doses. These parameters are associated with different factors representative of different microbial pathways of the fermentation. Therefore, at the end, we are able to identify which factors/pathways contribute the most to CH_4_ and VFA variation, and to quantify these contributions. This can be useful, in addition to *in vitro* and *in vivo* studies, to know on which factors to play to reduce CH_4_ production.

Also, the *in silico* or numerical experiment framework in which SA is conducted was used to analyze the uncertainty associated with the outputs of interest over time.

This work also addresses the limitations pointed out by Tedeschi (2021) in the evaluation process of the model in Muñoz-Tamayo et al. (2021), which did not include SA to assess the impact of model parameters on the model outputs.

### 2. Methods

First, a full description of the mechanistic model studied in this work is performed. This description includes the conceptual representation of the rumen fermentation phenomenon, the presentation of model equations, and the explanation of how the effect of AT on the fermentation was integrated in the model. Second, all the elements of the sensitivity analysis implementation are presented, with the description of the IPs studied, simulation scenarios considered and sensitivity analysis method implemented. Finally, we address how the uncertainty of model simulations were investigated.

### 2.1. Presentation of the mechanistic model

The model represents a rumen simulation technique (RUSITEC) system on a daily scale. Characteristics of the simulated RUSITEC system were taken from the setting used in Belanche et al., 2017. In the model, the rumen fermentation is represented as a reactor with a liquid phase of volume *V*_1_ and a gas phase of volume *V*_g_. The total volume of the system was set to 0.8 L with a separation of 0.74 L in liquid phase (*V*_1_) and 0.06 L in gas phase (*V*_g_). The feed “ingested” by the reactor is the input flux of the system. The biochemical mechanisms occurring during the degradation of the feed by the microbial community of the rumen with AT supply are represented according to assumptions described in Muñoz-Tamayo et al. (2016,2021). The model has 19 biochemical components in liquid and gas phase. The dynamics of the 19 state variables are represented by ordinary differentials equations. The model considers that system is completely mixed, which means that spatial variation is not directly modeled.

#### 2.1.1. Phenomena representation

The structure of the rumen fermentation model used in this study is determined by the representation of two phenomena namely the flow transport and microbial fermentation. The first is a biochemical (*e*.*g*., liquid-gas transfer) and physical phenomenon (*e*.*g*., output flow due to dilution rate) describing the transport fluxes in the system, here represented as a reactor. The second is a biological phenomenon describing the microbial fermentation of feeds.

The system studied is displayed in Figure 1.

**Figure 1.**
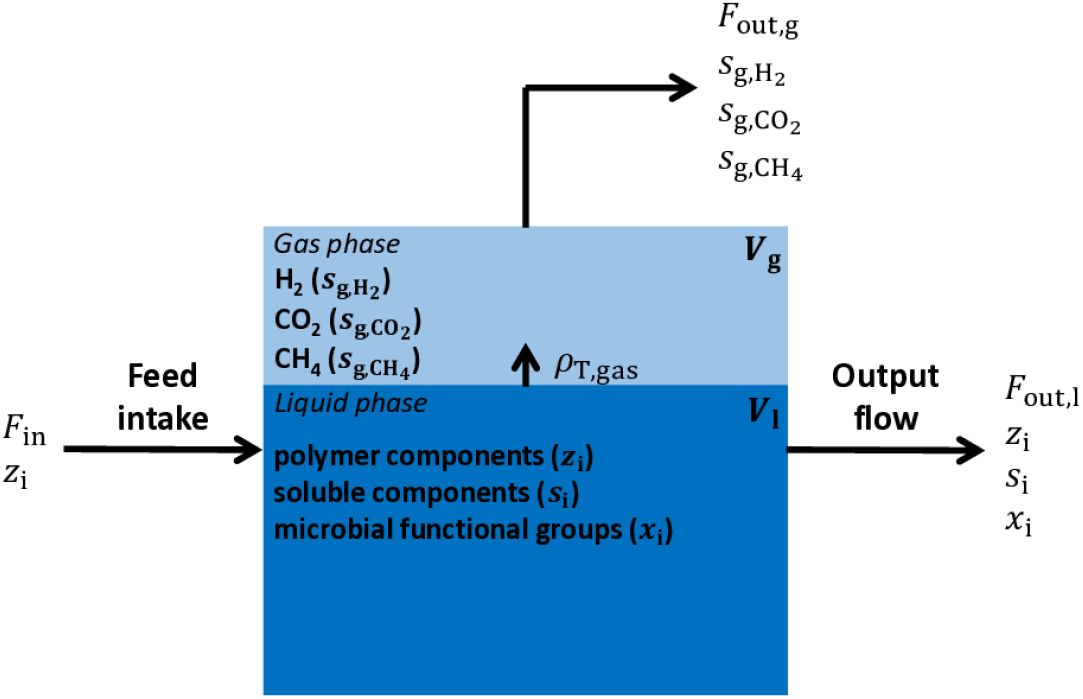
Representation of the *in vitro* continuous system. The system is composed of liquid and gas phases associated with the volumes *V*_1_ and *V*_g_, respectively. The liquid-gas transfer phenomena occur with the rate 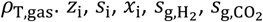 and 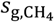 correspond to the concentrations of the biochemical components produced from the fermentation (the polymer components, the soluble components, the microbial functional groups, the hydrogen in gas phase, the carbon dioxide in gas phase and the methane in gas phase, respectively). They are the state variables of the Muñoz-Tamayo et al. (2021) model. *F*_in_ represents the input flux of the system, which is computed based on the feed intake described by the polymer component concentration (*z*_i_). *F*_out,1_ and *F*_out,g_ represent the output fluxes in liquid and gas phase, respectively.

This system is represented as a reactor system similar to engineering anaerobic digestion reactors (Batstone et al., 2002). It should be noted that Muñoz-Tamayo et al. (2021) model is a batch system, not considering input and output flow rates. The daily total dry matter ingested (*DM*_*total*_) was of 11.25 g. In our model, the dry matter intake (DMI) at a given time *t* was set as a dynamic equation determined by the number of feed distributions (*n*_*r*_), which corresponds to the act of the feed being distributed in the RUSITEC system. For the feed distribution *j*, the *DMI* kinetics follows

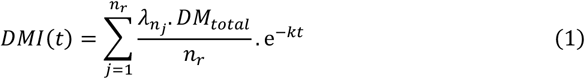

Where *DM*_*total*_ is the total quantity of dry matter (DM) ingested in one day (g), 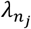 is the fraction of *DM*_*total*_ supplied in the distribution *j* and *k* (h^−1^) is the intake kinetic rate. *DM*_*total*_ was set to 11.25 g supplied in two feed distributions (*n*_*r*_ = 2). We set the first feed distribution to account for 70% of the total DM 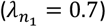. This configuration provides a DMI kinetics composed of two distributions with a significantly greater amount of DM ingested during the first intake and a medium intake kinetic (Figure 2).

**Figure 2.**
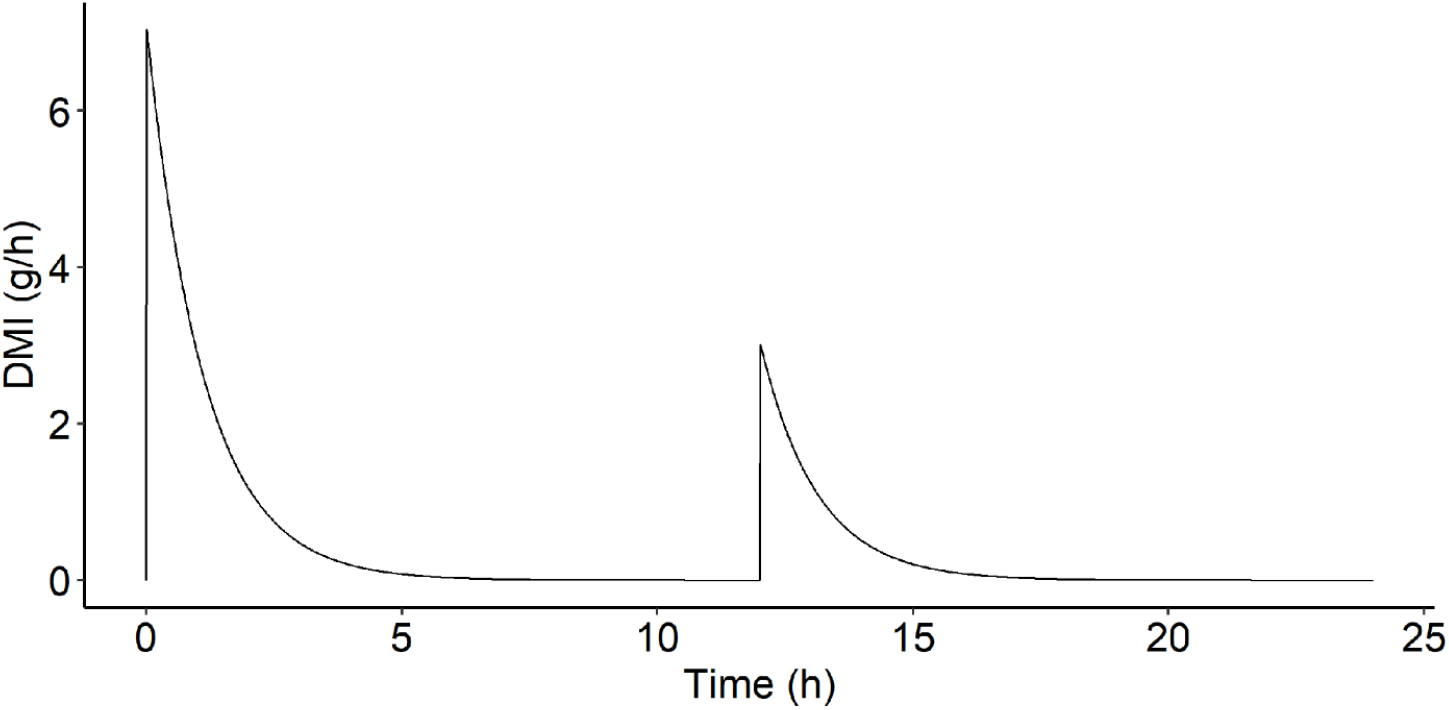
Dry matter intake (DMI, g/h) over time (h) simulated for one day with 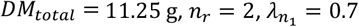 and k = 0.015.

The feed intake constitutes the input flux of the system. The feed is degraded by the rumen microbiota, leading to the production of several components in liquid and gas phase. Polymer components, soluble components and microbial functional groups are the components in liquid phase, and hydrogen, carbon dioxide and CH_4_ are the components in gas phase. Chemical compounds leave the system in liquid and gas phases as shown in Figure 1.

The representation of the fermentation process is displayed in Figure 3. It corresponds strictly to what happens in the liquid phase (blue box of Figure 1).

**Figure 3.**
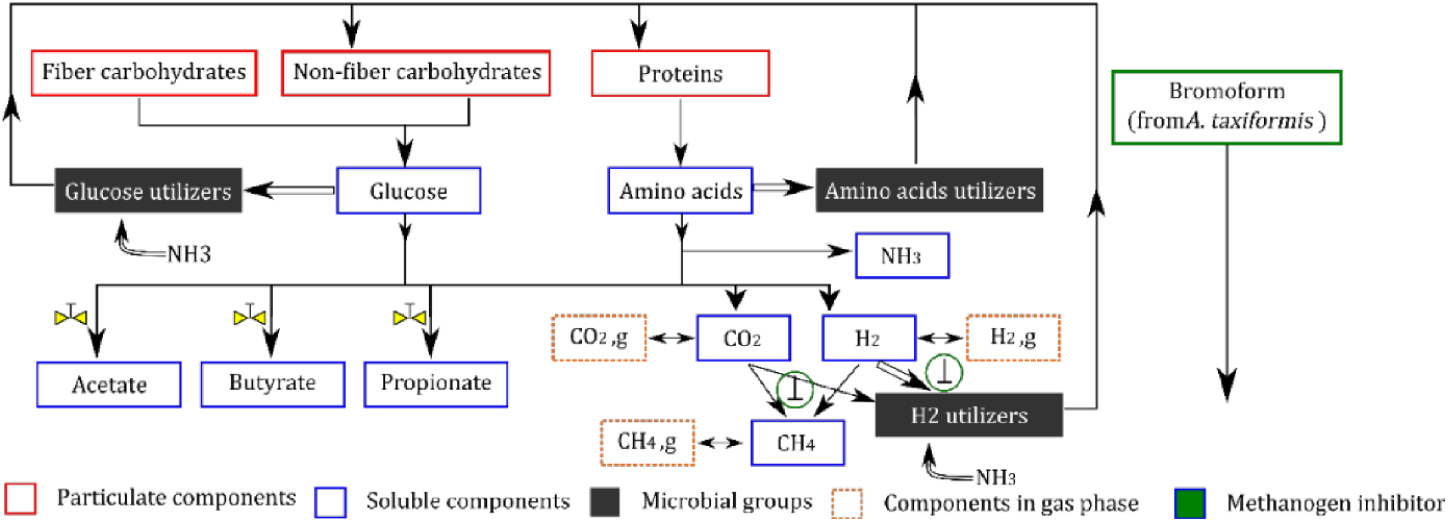
Representation of the *in vitro* rumen fermentation from Muñoz-Tamayo et al., 2021 model. This conceptual representation is based on biochemical assumptions described in Muñoz-Tamayo et al. (2016, 2011). The main assumptions are: 1) three polymer components are considered in the rumen: fiber carbohydrates, non-fiber carbohydrates and proteins, 2) hydrolysis of polymer components releases glucose (for fibers and non-fibers) and amino acids (for proteins), constituting two of the three soluble limiting substrates available in the rumen. The last soluble limiting substrate available is hydrogen, 3) the rumen microbiota is represented by three microbial functional groups (glucose utilizers, amino acids utilizers and hydrogen utilizers) determined by the microbial utilization of the three soluble limiting substrates in the fermentation pathway, 4) the utilization of the soluble substrates by biological pathways is done towards two mechanisms: product formation (single arrows) and microbial growth (double arrows), 5) acetate, propionate and butyrate are the only volatile fatty acids produced from the fermentation and 6) methane, carbon dioxide and hydrogen are the gas outputs of the fermentation. The inclusion of bromoform as the inhibitor compound of AT impacted the fermentation *via* two mechanisms. First, the bromoform has a direct inhibition of the growth rate of methanogens, resulting in a CH_4_ production reduction and hydrogen accumulation (represented by 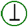). Second, the bromoform affects indirectly, through the hydrogen accumulation, the flux allocation towards VFA production, as hydrogen exerts control on this component (Mosey, 1983) (represented by 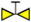).

Biochemical assumptions used to describe the fermentation and the effect of AT are detailed in Muñoz-Tamayo et al. (2016, 2021). The main ones are described in the caption of Figure 3. The resulting model comprises 19 state variables corresponding to 19 biochemical component concentrations in liquid and gas phases.

#### 2.1.2. System characteristics, intake and dietary scenarios

The consideration of feed intake (Figure 1) in the model studied allows to simulate dietary scenarios. In the system, the feed intake is composed of one intake scenario, which defines the intake dynamic, and one diet scenario, which defines the diet composition. The intake scenario is simulated before running the model by computing Equation 1 setting parameters with expert knowledge, conducting to Figure 2. Whereas, the diet scenario is set when computing the input flux *F*_*i*,in_ associated with the three polymer components (Equation 2, presented later). The fractions of fiber carbohydrates, non-fiber carbohydrates and proteins in the diet (*w*_ndf_, *w*_nsc_, *w*_pro_) were set using data from an *in vitro* study assessing several dietary CH_4_ mitigation strategies, including AT, on the fermentation (Chagas et al., 2019). This experiment tested the impact on CH_4_ production of several doses of AT from 0 to 1% DM (containing 6.84 mg/g dry weight of bromoform) in the diet. The diet was composed of 38.7% DM of neutral detergent fiber (*w*_ndf_), 39.7% DM of non-structural carbohydrates (*w*_nsc_) and 16% DM of crude proteins (*w*_pro_). We analyzed three simulation scenarios: A control treatment with 0% of AT, a low treatment with 0.25% of AT and a high treatment with 0.50% of AT (Figure 4).

**Figure 4.**
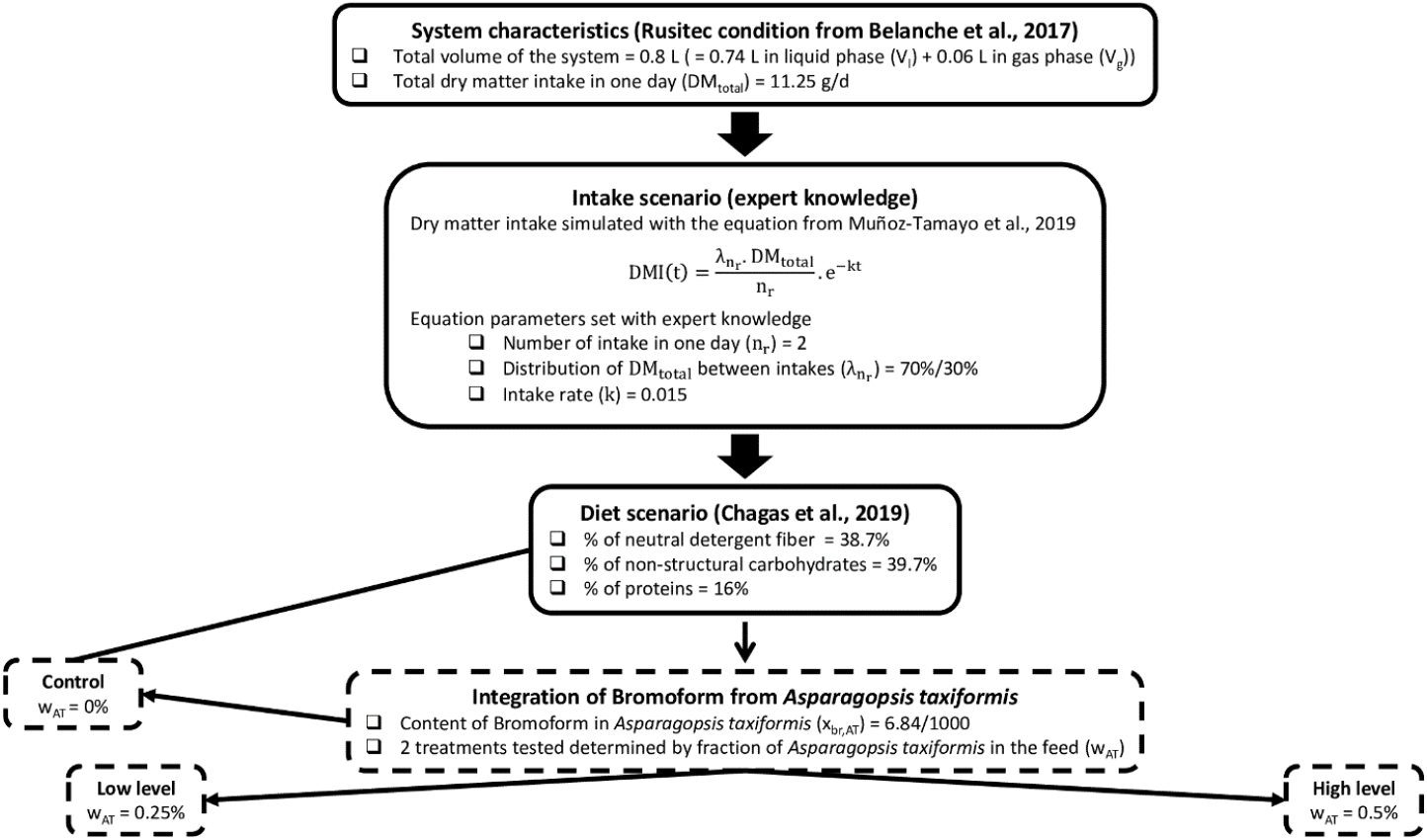
Summary of system characteristics, intake scenario and diet scenario simulated with the mechanistic model.

The initial condition of bromoform concentration was set to zero for all the three treatments. These three scenarios are representative of a typical forage-based diet for dairy or beef cattle, testing three reasonable doses of AT, which were also studied *in vivo* (Roque et al., 2021).

#### 2.1.3. Model equations

Model state variables are defined as **ξ** = (**z, s, x, s**_**g**_), where **z** = (*z*_ndf_, *z*_nsc_, *z*_pro_) is the vector of concentrations of the polymer components (neutral detergent fiber (*z*_ndf_), non-structural carbohydrates (*z*_nsc_) and proteins (*z*_pro_); g/L),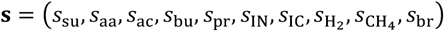 is the vector of concentrations of the soluble components (sugars (*s*_su_), amino acids (*s*_aa_), acetate (*s*_ac_), butyrate (*s*_bu_), propionate (*s*_pr_), inorganic nitrogen (*s*_IN_), inorganic carbon (*s*_IC_), hydrogen 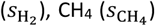 and bromoform (*s*_br_); mol/L), 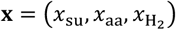 is the vector of concentrations of the microbial functional groups (sugars utilizers (*x*_su_), amino acids utilizers (*x*_aa_) and hydrogen utilizers 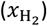; mol/L), 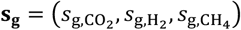 is the vector of concentrations in gas phase (carbon dioxide 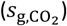, hydrogen 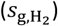 and 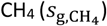; mol/L). The polymer components include an input flux in their equation and all the components are associated with an output flux in liquid or gas phase. The input flux (g/(Lh)) of polymer components is described as

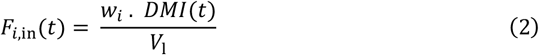

where *w*_*i*_ is the fraction of polymer component *i* in the diet of the animal, *DMI* is the DM intake (g/h) with a total DM of 11.25 g split in two feed distributions along the day with the first distribution accounts for 70% of the total DM, and *V*_1_ the volume in liquid phase of the rumen (L).

The output flux in liquid phase (g/(Lh) for polymer components and mol/(Lh) for soluble and microbial functional groups components) is described as

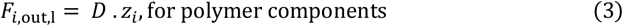

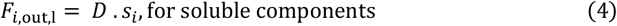

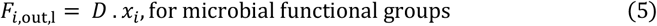

where *D* is the dilution rate (*D* = 0.035 h^−1^, Bayat et al., 2011), *z*_*i*_ is the concentration of polymer component *i, s*_*i*_ is the concentration of soluble component *i* and *x*_*i*_ is the concentration of microbial functional group *i*.

The output flux in gas phase (mol/(Lh)) is described as

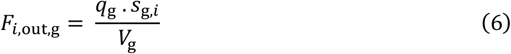

where 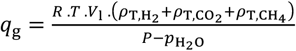 is the output flow of gas phase (L/h) wit *R* the ideal gas constant (barL/(molK)), *T* the temperature of the rumen 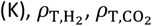 and 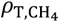 the liquid-gas transfer phenomena rates of hydrogen, carbon dioxide and CH_4_ (mol/(Lh)), respectively, *P* the total pressure (bars) and 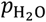 the partial pressure of water vapor (bar). *s*_g,*i*_ is the concentration of component *i* in gas phase (mol/L) and *V*_g_ is the volume in gas phase of the rumen (L).

Model equations are derived from mass balance equations described below.

##### For polymer components

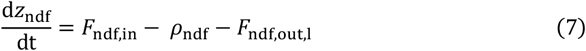

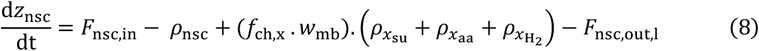

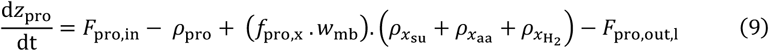

where *F*_ndf,in_, *F*_nsc,in_ and *F*_pro,in_ are the input fluxes of neutral detergent fiber, non-structural carbohydrates and proteins (g/(Lh)), respectively. *ρ*_ndf_, *ρ*_nsc_ and *ρ*_pro_ are the hydrolysis rate functions of polymer components (g/(Lh)), indicating the kinetic of hydrolysis of polymer components. These functions are described as

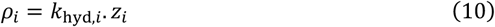

With *k*_hyd,*i*_ the hydrolysis rate constant (h^−1^) and *z*_*i*_ the concentration of polymer component *i* (g/L). *F*_ndf,out_, *F*_nsc,out_ and *F*_pro,out_ are the output fluxes of polymer components (g/(Lh)). The middle part of equations (8) and (9) represents the recycling of dead microbial cells where *f*_ch,x_, *f*_pro,x_ are the fractions of carbohydrates and proteins of the biomass, *w*_mb_ is the molecular weight of microbial cells (g/mol) and 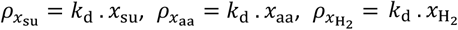 are the cell death rate of sugars utilizers, amino acids utilizers and hydrogen utilizers (mol/(Lh)) with *k*_d_ the rate of dead of microbial cells (h^−1^).

##### For soluble components

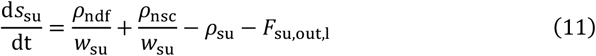

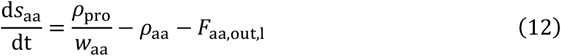

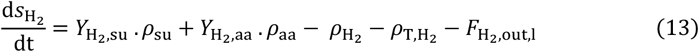

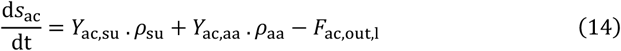

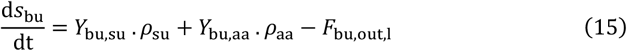

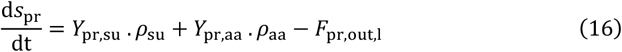

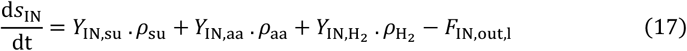

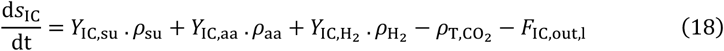

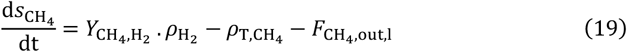

Let us detail how the amino acids are produced and used by the biological pathways in the fermentation process. Equation (12) indicates that amino acids are produced (positive sign in the equation) from the degradation of proteins, occurring with the kinetic rate *ρ*_pro_ (g/L h), which is divided by the molecular weight of the amino acids *w*_aa_ (g/mol). Moreover, amino acids are utilized (negative sign in the equation) by the biological pathways during the fermentation with the kinetic rate *ρ*_aa_ (mol/Lh). The kinetic rate *ρ*_aa_ is a function indicating the kinetic of utilization of amino acids during the fermentation and is described as

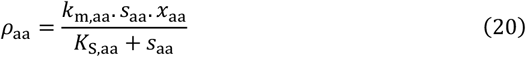

With *k*_m,aa_ the maximum specific utilization rate constant of amino acids (mol substrate/(mol biomass h)), *s*_aa_ the concentration of amino acids (mol/L), *x*_aa_ the concentration of amino acids-utilizing microbes (mol/L) and *K*_S,aa_ the Monod affinity constant associated with the utilization of amino acids (mol/L). *F*_aa,out_ is the output flux of amino acids concentration (mol/(Lh)). Then, further in the fermentation, amino acids are utilized by the specific microbial functional group *x*_aa_ and contributed to the production of hydrogen (Equation 13), VFA (Equations 14, 15, 16), inorganic nitrogen (Equation 17) and inorganic carbon (Equation 18) with a stoichiometry represented by the yield factors 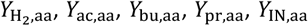 and *Y*_IC,aa_, respectively. These components are also produced from glucose metabolism. In the last step of the biochemical conversion cascade, inorganic nitrogen and inorganic carbon are utilized during the reaction of hydrogen utilization in liquid phase with the kinetic rate function 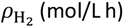, described similarly as Equation (20). An additional term (*I*_br_) is included to represent the inhibition effect of bromoform on the hydrogen utilizers (methanogens) as detailed later on. Hydrogen in liquid phase is also associated with a transfer phenomenon with hydrogen in gas phase given by the rate 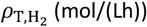). This liquid-gas transfer phenomenon also concerns carbon dioxide with the rate 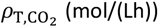) and CH_4_ (Equation 19) with the rate 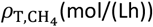). The general equation of the liquid-gas transfer rate is described as

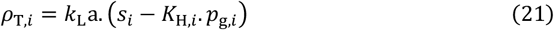

With *k*_L_ a the mass transfer coefficient (h^−1^), *s*_*i*_ the concentration (mol/L), *K*_H,*i*_ the Henry’s law coefficient (M/bar) and *p*_g,*i*_ the partial pressure (bars) of soluble component *i*.

Finally, CH_4_ in liquid phase is produced using hydrogen in liquid phase with the stoichiometry 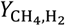.

##### For microbial functional groups

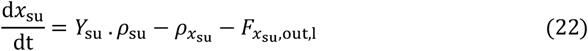

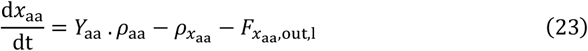

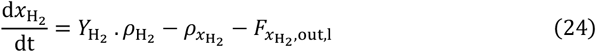

Microbial functional groups of glucose utilizers (Equation 22), amino acid utilizers (Equation 23) and hydrogen utilizers (Equation 24) are produced from their respective substrates with the yield factors *Y*_su_, *Y*_aa_ and 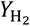, respectively.

##### For the gas phase

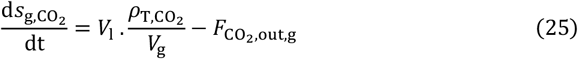

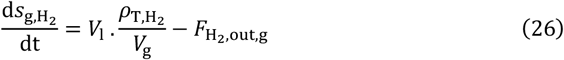

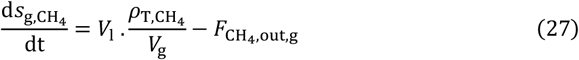

The dynamics of carbon dioxide (Equation 25), hydrogen (Equation 26) and CH_4_ (Equation 27) in gas phase are driven by the liquid-gas transfer phenomena given by the rates 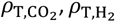 and 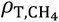 (mol/(Lh)), respectively.

Model parameters were either set with values extracted from the literature (Batstone et al., 2002; Serment et al., 2016), set with values reported from *in vitro* study providing the experimental data (Chagas et al., 2019) or estimated using the maximum likelihood estimator as reported in Muñoz-Tamayo et al. (2021). In the present study, initial conditions of state variables were determined by running the model for 50 days without AT supply (control condition). The idea was to reach a quasi-steady state of the state variables. Values corresponding to the last time step simulated were selected as initial conditions of the model for the further analysis explained below.

#### 2.1.4. Integration of the macroalgae *Asparagosis taxiformis*

The integration of bromoform contained in AT conducted to the incorporation of the 19^th^ state variable representing the dynamic of bromoform concentration.

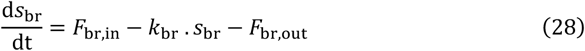

Where 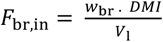 is the input flux of bromoform concentration (g/(Lh)) with *w*_br_ the fraction of bromoform in the diet of the animal. *k*_br_ corresponds to the kinetic rate of bromoform utilization (h^−1^) and *F*_br,out_ = *D* . *s*_br_ is the output flux of bromoform concentration (g/(Lh)). The value of *k*_br_ was obtained from data reported in Romero et al. (2023b).

The direct effect of bromoform on the CH_4_ production is represented through the factor *I*_br_ (Equation 29) impacting the kinetic rate function of hydrogen utilization 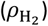. This factor is a function of the bromoform concentration and is modeled by a sigmoid shape. Whereas the indirect effect of bromoform on the flux allocation towards VFA production is represented through the flux allocation parameters *λ*, describing the three reactions driving flux allocation from glucose utilization to VFA production. *λ*_1_, *λ*_2_ and *λ*_3_ indicates the molar fraction of glucose utilized to produce acetate, to produce propionate and to produce butyrate, respectively. They follow *λ*_1_ + *λ*_2_ + *λ*_3_ = 1. λ_1_ (Equation 30) and *λ*_2_ (Equation 31) are represented by affine functions described below.

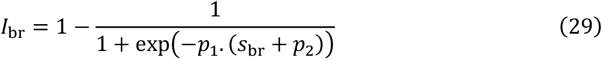

With *s*_br_ the bromoform concentration (g/L) and *p*_1_, *p*_2_ the parameters of sigmoid function.

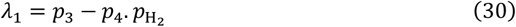

With 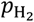 the hydrogen partial pressure (bars) and *p*_3_, *p*_4_ the parameters of affine function.

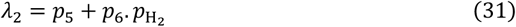

With 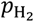 the hydrogen partial pressure (bars) and *p*_5_, *p*_6_ the parameters of affine function. These factors are displayed in Figure 5.

**Figure 5.**
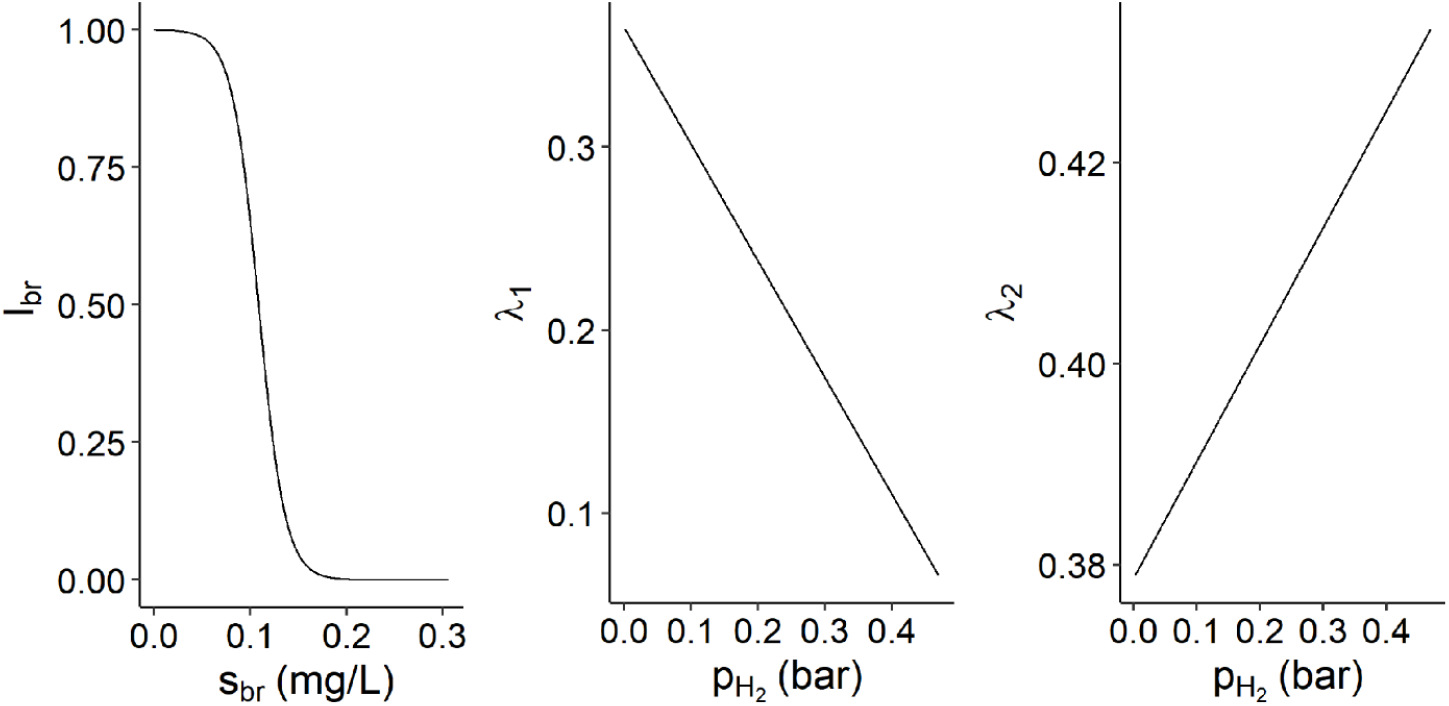
Representation of the direct effect of *Asparagosis taxiformis* on the methane production (*I*_br_) against the bromoform concentration (*s*_br_, mg/L), and of the indirect effect of *Asparagosis taxiformis*, through the hydrogen accumulation, on the flux allocation towards acetate production (*λ*_1_) and propionate production (*λ*_2_) against the hydrogen partial pressure 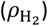.

It should be noted that the model version of Muñoz-Tamayo et al. (2021) includes an inhibition factor of glucose utilization by hydrogen 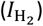. This factor was incorporated to account for the reduced production of VFA under AT supply observed in the experiments of Chagas et al. (2019). However, in the present study, we decided not to include this term. Some studies have shown that high doses of AT decrease total VFA both *in vitro* (Chagas et al., 2019; Kinley et al., 2016; Machado et al., 2016; Terry et al., 2023) and *in vivo* (Li et al., 2016; Stefenoni et al., 2021). However, in other studies the total VFA was unaffected by AT supplementation under *in vitro* (Romero et al., 2023b; Roque et al., 2019) and *in vivo* (Kinley et al., 2020) conditions. These discrepancies might be due to the variability of physical processing of the macroalgae (*e*.*g*., drying, storage). The incorporation of the hydrogen inhibition factor was indeed challenged by Henk Van Lingen in the evaluation of the model in Muñoz-Tamayo et al., (2021) (Tedeschi, 2021). Accordingly, we acknowledge that this aspect requires further studies, and it is not then included in the present work. We then run again the calibration of the model without the 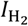 factor under batch conditions using the experimental data from Chagas et al. (2019) to estimate the parameters from equations (28-30). In the process, with the aim of model simplification, we set the allocation factors *λ*_1_, *λ*_2_ as linear functions of 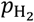. The updated version of the model under batch conditions is available at Muñoz-Tamayo (2020).

### 2.2. Sensitivity analysis

Based on the representation of Figure 3, we focus on the hydrolysis of polymer components (fiber carbohydrates, non-fiber carbohydrates and proteins), the fermentation of microbial functional groups (glucose utilizers, amino acids utilizers and hydrogen utilizers) and the effect of bromoform on rumen fermentation. They are the components of interest, those whose impact on the variability of CH_4_ and VFA production is to be studied, when implementing the SA.

We implemented a SA method for quantifying the contribution of 16 IPs to the variability of four state variables of the mechanistic model described in the previous section. This method is grounded on a strong theoretical framework and provide easy-to-interpret sensitivity indices (SI). Moreover, the SI were computed over time allowing studying the dynamics of IP sensitivity during the fermentation. The Shapley effects (Owen, 2014) were computed for quantifying the individual, interaction and dependence/correlation effects of an IP to the variability of an output. Although the model studied here provides no correlated or dependent IPs, this method was used to introduce a new SA method in the animal nutrition field. In addition to Shapley effects, we explored also the Sobol method. The interested reader is referred to the supplementary material (Blondiaux et al., 2024).

This work was done with the support of MESO@LR-Platform at the University of Montpellier, which was used to run the algorithms computing Shapley effects. They were run using one node, 28 cores and 125 GB of RAM of memory.

#### 2.2.1. Input parameters and output variables studied

State variables of the model considered as output variables of the SA were the rate of CH_4_ production (in gas phase) 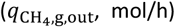 and VFA (acetate (*s*_ac_), butyrate (*s*_bu_) and propionate (*s*_pr_)) concentrations (mol/L). The model was run for four days, similarly to the RUSITEC of Roque et al., (2019), and the SA was performed on the last day of simulation, from 72 to 96h with a time step of 1h.

Components constituting the model represent different factors associated to microbial pathways involved in *in vitro* rumen fermentation. These factors include polymer hydrolysis and microbial growth. The sensibility of hydrolysis rate constants associated with the three polymer components (*k*_hyd,ndf_, *k*_hyd,nsc_ and *k*_hyd,pro_, h^−1^) was studied for quantifying the impact of feed polymer hydrolysis on output variables of interest during the fermentation. In addition, the sensibility of maximum specific utilization rate constants (*k*_m,su_, *k*_m,aa_ and 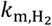, mol substrate/(mol biomass h)) and Monod constants (*K*_S,su_, *K*_S,aa_ and 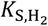, mol/L) associated with the three microbial groups was studied for quantifying the impact of microbial growth on output variables of interest during the fermentation.

In addition to the IPs quantifying the impact of polymer components and microbial functional groups, IPs related to the effect of bromoform on the fermentation were considered. The kinetic rate constant of bromoform utilization (*k*_br_, h^−1^), quantifying the consumption of anti-methanogenic compounds, was added to the SA. Moreover, the parameters of sigmoid and affine functions associated with the factor representing the impact of bromoform on methanogens (*I*_br_ associated with parameters *p*_1_ and *p*_2_) and with the flux allocation from glucose utilization to VFA production (*λ*_1_ associated with parameters *p*_3_ and *p*_4_, and *λ*_2_ associated with parameters *p*_5_ and *p*_6_) were added to the SA. Therefore, in total, 16 IPs were considered.

The first step in SA was to set the variability space of IPs. To perform that, information about the variability of each IP was required. This information was available from two sources: data and expert knowledge (Table 3). Based on the low number of data available for each IP, uniform distributions were selected for quantifying hydrolysis rate constants, maximum specific utilization rate constants and Monod constants variability. Lower and upper bounds of uniform distributions were set by selecting the minimum and maximum values among all the references. Furthermore, the parameters associated with the effect of bromoform on the fermentation were not biological parameters and no data were available for modeling their variability. Therefore, a uniform distribution varying of ± 10% the baseline model parameter value was used for parameters *p*_1_ to *p*_6_.

**Table 1.**
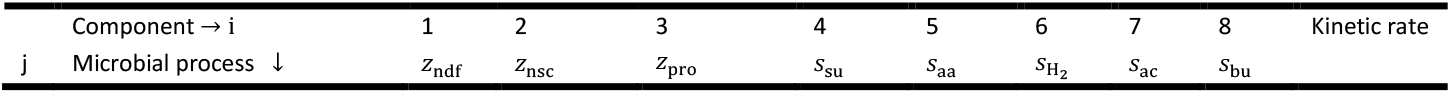
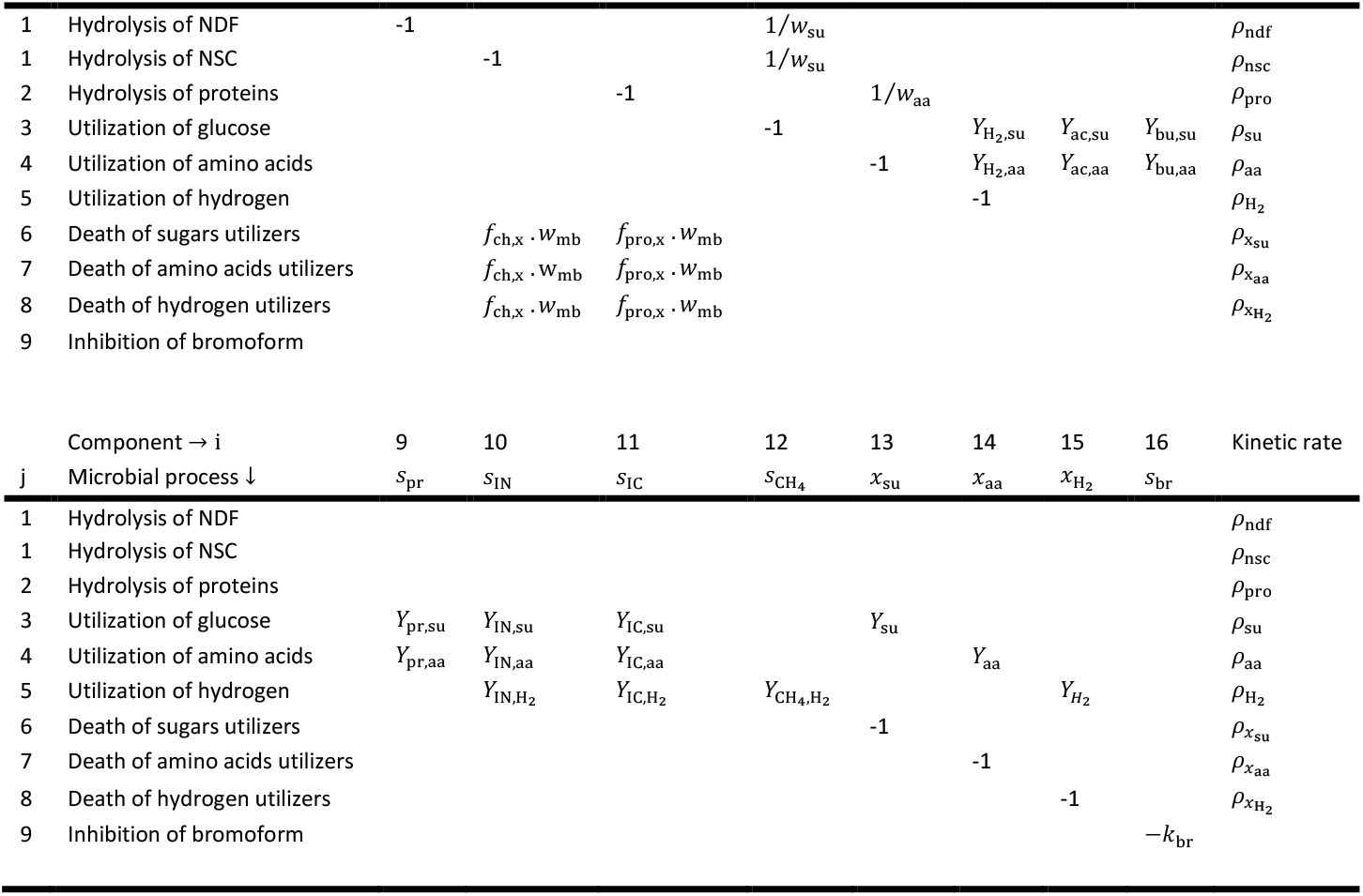
Stoichiometry matrix of biochemical reactions occurring during the rumen fermentation.

| Component → i | 1 | 2 | 3 | 4 | 5 | 6 | 7 | 8 | Kinetic rate |
| --- | --- | --- | --- | --- | --- | --- | --- | --- | --- |
| j Microbial process ↓ | $z_{ndf}$ | $z_{nsc}$ | $z_{pro}$ | $s_{su}$ | $s_{aa}$ | $s_{H_2}$ | $s_{ac}$ | $s_{bu}$ | |

|  |  |  |  |  |  |  |  |  |  |  |
| --- | --- | --- | --- | --- | --- | --- | --- | --- | --- | --- |
| 1 | Hydrolysis of NDF | -1 | | | $1/w_{su}$ | | | | | $\rho_{ndf}$ |
| 1 | Hydrolysis of NSC | | -1 | | $1/w_{su}$ | | | | | $\rho_{nsc}$ |
| 2 | Hydrolysis of proteins | | | -1 | | $1/w_{aa}$ | | | | $\rho_{pro}$ |
| 3 | Utilization of glucose | | | | -1 | | $Y_{H_2,su}$ | $Y_{ac,su}$ | $Y_{bu,su}$ | $\rho_{su}$ |
| 4 | Utilization of amino acids | | | | | -1 | $Y_{H_2,aa}$ | $Y_{ac,aa}$ | $Y_{bu,aa}$ | $\rho_{aa}$ |
| 5 | Utilization of hydrogen | | | | | | -1 | | | $\rho_{H_2}$ |
| 6 | Death of sugars utilizers | | $f_{ch,x} \cdot w_{mb}$ | $f_{pro,x} \cdot w_{mb}$ | | | | | | $\rho_{x_{su}}$ |
| 7 | Death of amino acids utilizers | | $f_{ch,x} \cdot w_{mb}$ | $f_{pro,x} \cdot w_{mb}$ | | | | | | $\rho_{x_{aa}}$ |
| 8 | Death of hydrogen utilizers | | $f_{ch,x} \cdot w_{mb}$ | $f_{pro,x} \cdot w_{mb}$ | | | | | | $\rho_{x_{H_2}}$ |
| 9 | Inhibition of bromoform |  |  |  |  |  |  |  |  |  |
| | Component → i | 9 | 10 | 11 | 12 | 13 | 14 | 15 | 16 | Kinetic rate |
| j | Microbial process ↓ | $s_{pr}$ | $s_{IN}$ | $s_{IC}$ | $s_{CH_4}$ | $x_{su}$ | $x_{aa}$ | $x_{H_2}$ | $s_{br}$ | |
| 1 | Hydrolysis of NDF | | | | | | | | | $\rho_{ndf}$ |
| 1 | Hydrolysis of NSC | | | | | | | | | $\rho_{nsc}$ |
| 2 | Hydrolysis of proteins | | | | | | | | | $\rho_{pro}$ |
| 3 | Utilization of glucose | $Y_{pr,su}$ | $Y_{IN,su}$ | $Y_{IC,su}$ | | $Y_{su}$ | | | | $\rho_{su}$ |
| 4 | Utilization of amino acids | $Y_{pr,aa}$ | $Y_{IN,aa}$ | $Y_{IC,aa}$ | | | $Y_{aa}$ | | | $\rho_{aa}$ |
| 5 | Utilization of hydrogen | | $Y_{IN,H_2}$ | $Y_{IC,H_2}$ | $Y_{CH_4,H_2}$ | | | $Y_{H_2}$ | | $\rho_{H_2}$ |
| 6 | Death of sugars utilizers | | | | | -1 | | | | $\rho_{x_{su}}$ |
| 7 | Death of amino acids utilizers | | | | | | -1 | | | $\rho_{x_{aa}}$ |
| 8 | Death of hydrogen utilizers | | | | | | | -1 | | $\rho_{x_{H_2}}$ |
| 9 | Inhibition of bromoform | | | | | | | | $-k_{br}$ | |

**Table 2.**
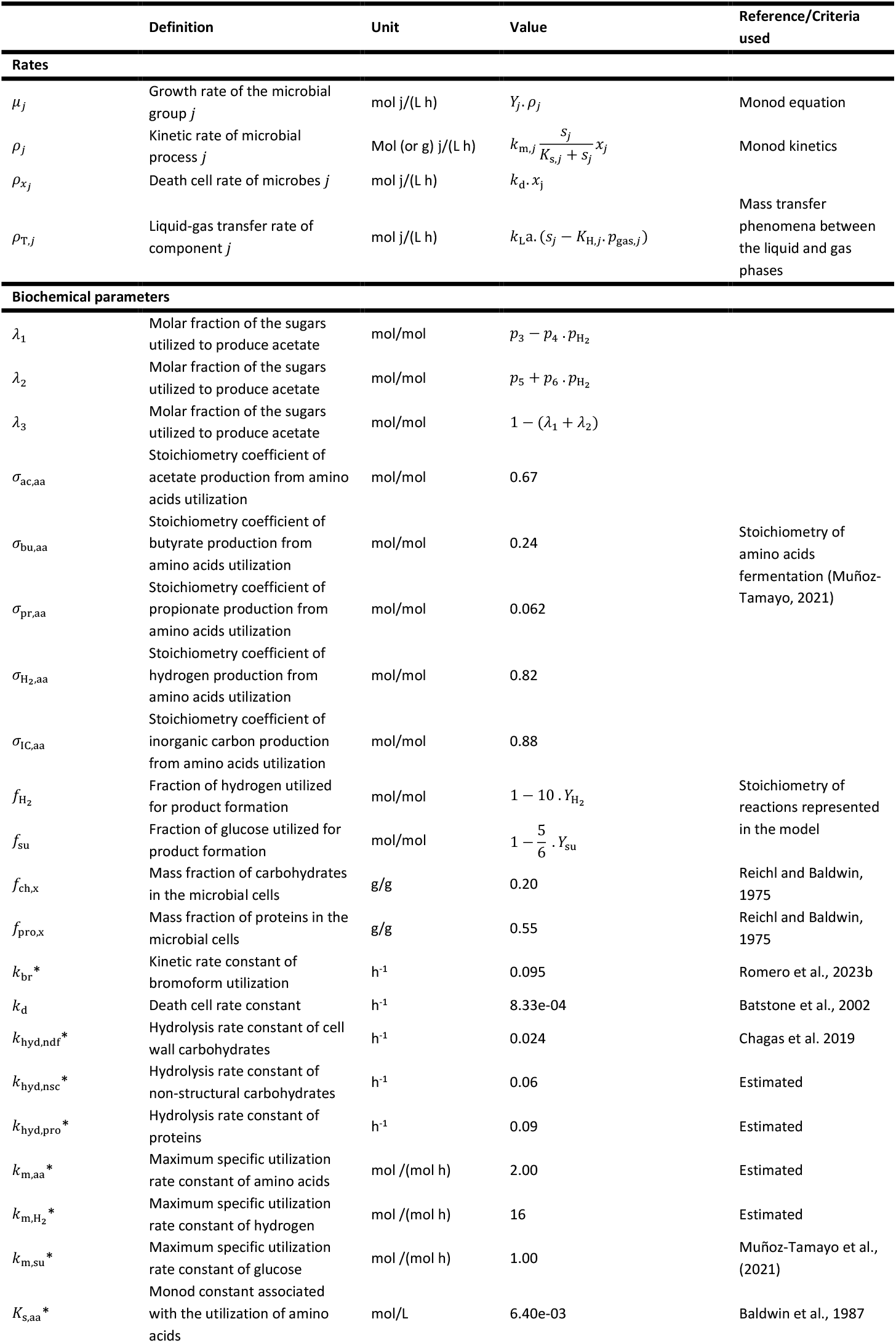
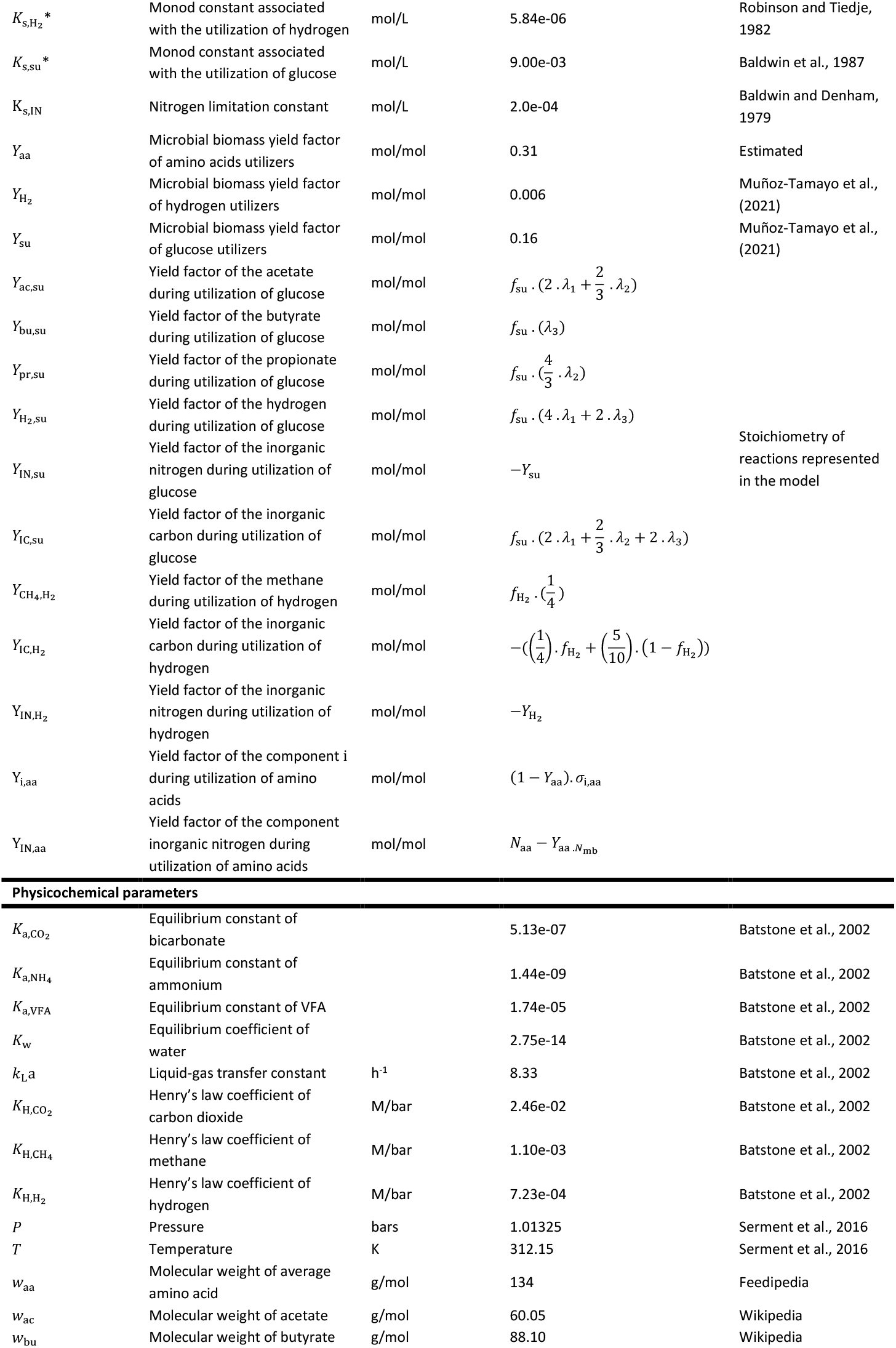
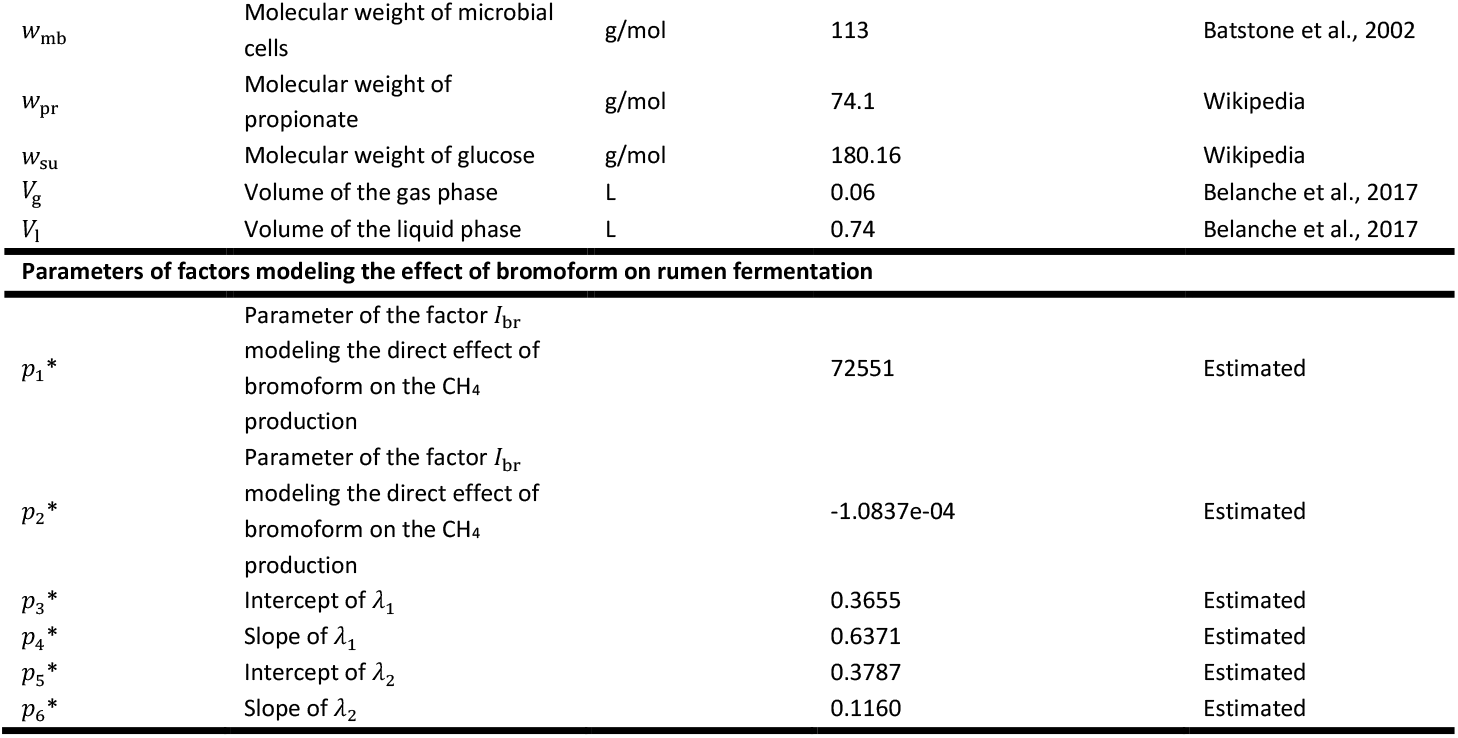
Model parameters. Parameters studied in the sensitivity analysis are highlighted with *.

**Table 3.**
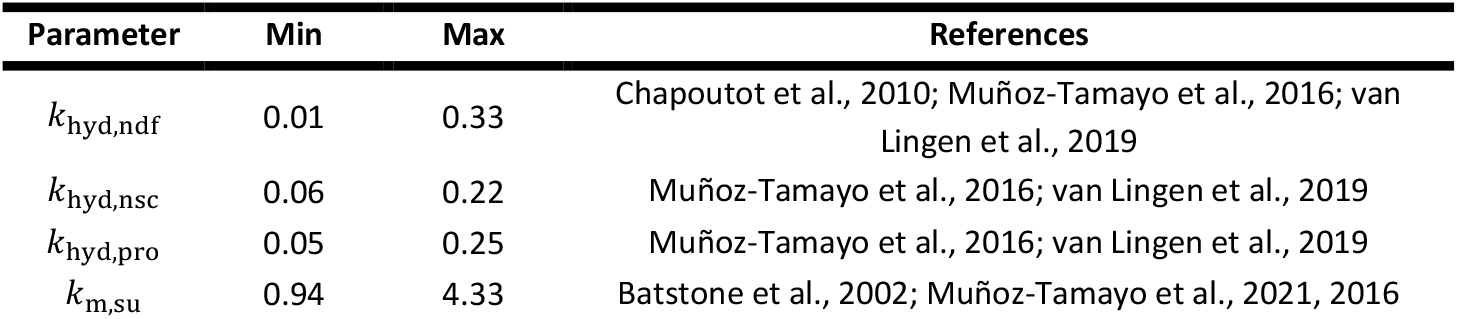
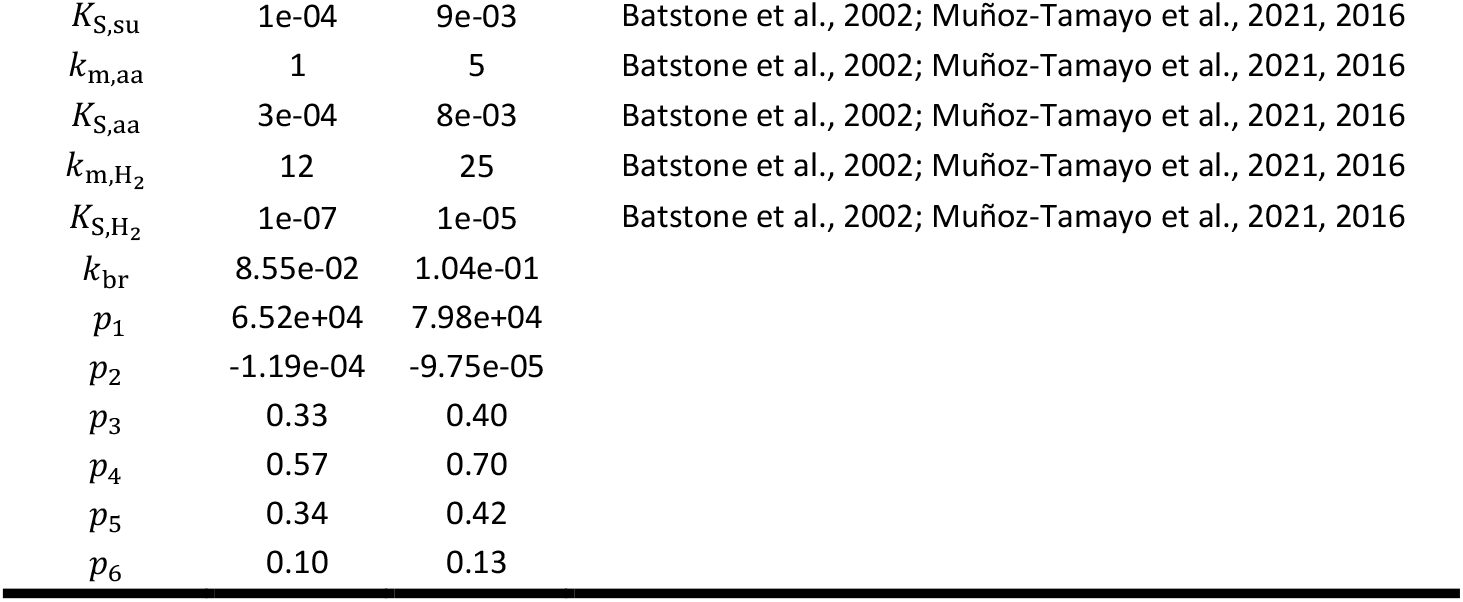
Variation range (minimum (min) and maximum (max)) and sources/references of uniform distributions used for exploring the variability of input parameters studied in the sensitivity analysis.

#### 2.2.2. Shapley effects

##### 2.2.2.1. Definition

The SA method implemented was the Shapley effects, which come from the field of cooperative game theory (Shapley, 1953). The Shapley effect of an IP *x*_i_ (*sh*_i_) measures the part of variability of the output variable caused by the variability of *x*_i_, and allocate to *x*_i_ a fair value regarding its individual contribution, its contribution due to interactions with other IPs and its contribution due to dependence/correlation with other IPs (Owen, 2014, Song et al., 2016). It is described as

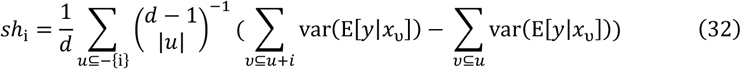

where *d* is the number of IPs, *u* ⊆ {1, … *d*] is a subset of IPs, *y* is the output variable and *x* is an IP.

##### 2.2.2.2. Interpretation

The Shapley effects are condensed and easy-to-interpret. Their sum is equal to 1, allowing us to interpret them as the percentage of contribution of the IPs to output variability. Nevertheless, the distinction of individual, interaction and dependence/correlation effects are not possible. Each IP is associated with one value, integrating the three effects.

##### 2.2.2.3. Numerical computation

Several methods are available for estimating the Shapley effects. In our study, the random permutation method was used (Song et al., 2016). This method provides a consistent estimation of the Shapley effects adapted in the case of numerous IPs (Iooss and Prieur, 2019). It is based on an alternative definition of the Shapley effects, expressing it in terms of all the possible IPs permutations (Castro et al., 2009). The computational cost of this method is *N*_*υ*_ + (*d* − 1)*N*_*o*_*N*_*i*_ with *N*_*υ*_ the sample size for estimating the output variance, *m* the number of permutations randomly sampled from the space of all the possible IP permutations, *d* the number of IPs considered, *N*_*o*_ the sample size for estimating the expectation and *N*_*i*_ the sample size for estimating the conditional variance. *N*_*o*_ and *N*_*i*_ were set at 1 and 3, respectively, as recommended in Song et al. (2016). In addition, *N*_*υ*_ = 1e04 and *m* = 1e04 were considered, conducting to 460000 model evaluations. Estimation of the Shapley effects was performed using the R package “sensitivity” (Iooss et al., 2023).

### 2.3. Uncertainty analysis

SA provides a framework combining an IP sampling matrix, developed by randomly drawing values from IP probability distributions (Table 3), to simulations of our four outputs of interest. This *in silico* framework was used for analyzing uncertainty associated with the simulations of CH_4_, acetate, butyrate and propionate concentrations (mol/L). Similarly to SA, uncertainty associated with outputs of interest was studied dynamically by computing summary statistics (median, standard deviation (SD), and quantiles 10 and 90%) and the coefficient of variation (CV) of the output simulations at each time step.

## 3. Results and discussion

### 3.1. Analysis of the simulations of the mechanistic model

#### 3.1.1. Comparison with *in vitro* and *in vivo* studies for methane production

Figure 6 displays the dynamic of 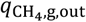 of the three dietary scenarios (control: 0% of AT, low treatment: 0.25% of AT and high treatment: 0.50% of AT) for a four days simulation.

**Figure 6.**
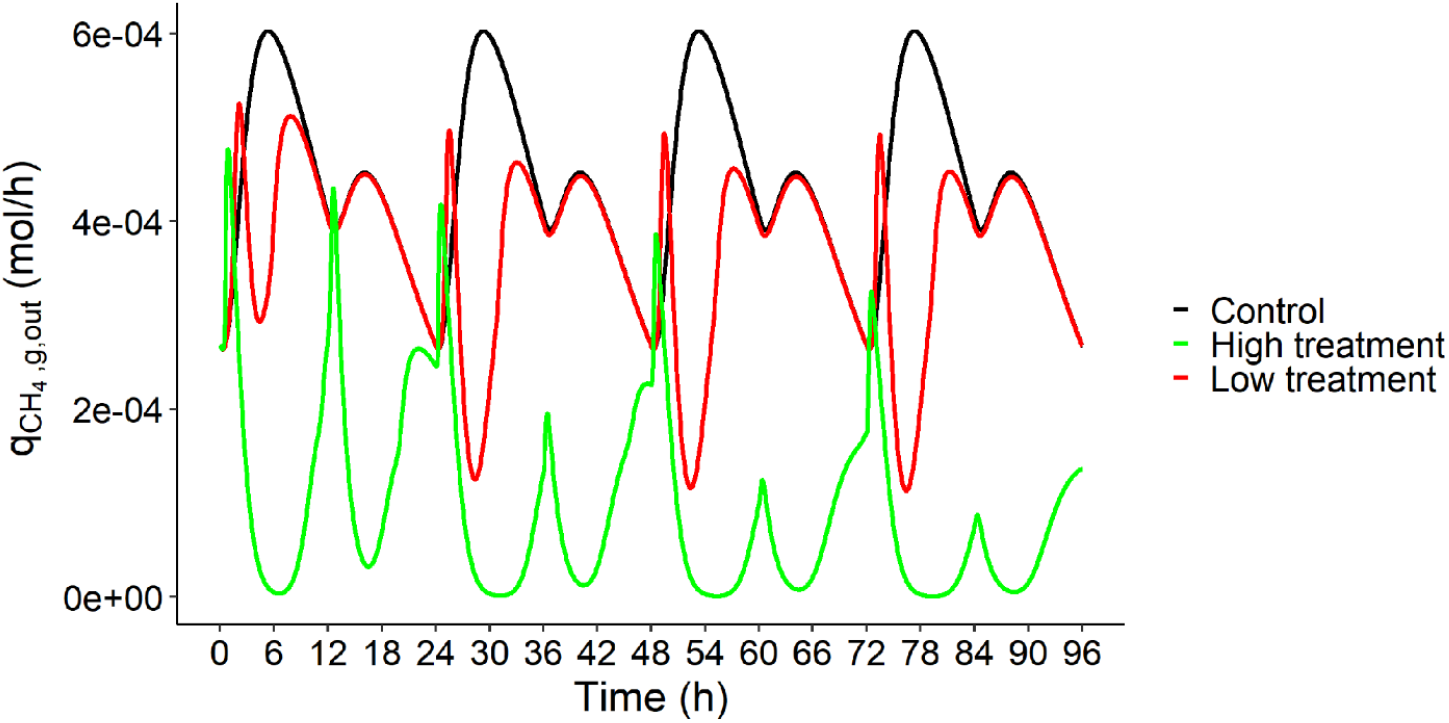
Rate of CH_4_ production in gas phase (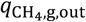, mol/h) over time (h) of the three dietary scenarios (control: 0% of *Asparagopsis taxiformis*, low treatment: 0.25% of *Asparagopsis taxiformis* and high treatment: 0.50% of *Asparagopsis taxiformis*) for a four days simulation.

Increasing the dose of AT decreased 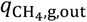 with, at the end of the four days of simulations, a CH_4_ (g/d) reduction of 17% between control and low AT treatment and of 78% between control and high AT treatment. This reduction increased from one day to the next with computed reductions of 9%, 14%, 16% and 17% between control and low AT treatment, and of 65%, 72%, 75% and 78% between control and high AT treatment, from day 1 to 4, respectively.

These reductions between AT treatments were lower than those reported in *in vitro* and *in vivo* studies. Chagas et al. (2019) indicated that the inclusion of AT (10 g/kg OM) decreased predicted *in vivo* CH_4_ production (mL/g DM) of 99% under *in vitro* condition. Whereas, the RUSITEC of Roque et al. (2019) and the *in vitro* study of Romero et al. (2023b) reported reductions of CH_4_ production (mL/g OM and mL, respectively) of 95% (with a 5% OM dose) and 97% (with a 2% DM dose), respectively.

Under *in vivo* conditions, Roque et al. (2021) tested AT doses similar to our simulations. This work reported *in vivo* CH_4_ production (g/d) reduction of 32.7 and 51.9% between control, and low and high AT treatments, respectively. The model simulated a lower reduction for low AT treatment and a higher reduction for high AT treatment.

These results highlighted that some interactions occurring during the fermentation are not represented in the model (e.g. forage wall content might inhibit the effect of AT). Improving the model involves a finer representation of the interactions between feed characteristics and fermentation, as discussed by Bannink et al. (2016).

#### 3.1.2. Analysis of the behavior of VFA proportions

The dynamic of VFA proportions and propionate to acetate ratio of the three dietary scenarios for a four days simulation is displayed in Figure 7.

**Figure 7.**
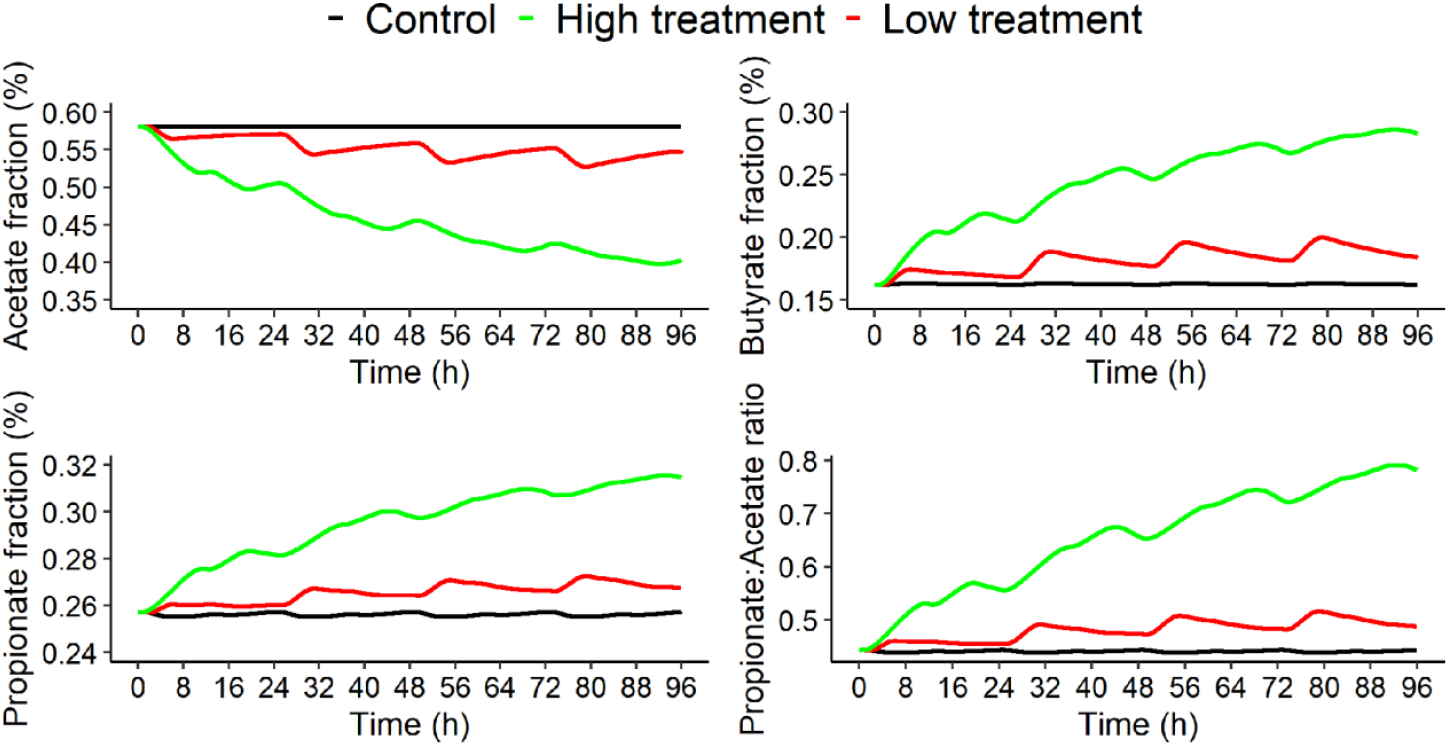
Acetate proportion (%), butyrate proportion (%), propionate proportion (%) and propionate to acetate ratio over time (h) of the three dietary scenarios (control: 0% of *Asparagopsis taxiformis*, low treatment: 0.25% of *Asparagopsis taxiformis* and high treatment: 0.50% of *Asparagopsis taxiformis*) for a four days simulation.

The dynamic of VFA proportions showed that increasing AT dose in the diet decreased acetate proportion of 5% and 31% at the end of the fermentation between control, and low and high AT treatments, respectively. Whereas, butyrate and propionate proportions increased when increasing AT dose in the diet, with increases at the end of the fermentation (t = 96h) of 13% and 74% for butyrate proportion and of 4% and 22% for propionate proportion between control, and low and high AT treatments, respectively. These behaviors were similar to those of *in vitro* studies. In Chagas et al. (2019), the AT treatment was associated with lower molar proportion of acetate (−75%) and higher molar proportions of propionate and butyrate (+ 38% and + 47%, respectively).

Propionate to acetate ratio was also associated with an increase of 10% between control and low AT treatment and of 76% between control and high AT treatment. This increase was highlighted in Roque et al. (2019) and Romero et al. (2023b).

These results highlight that the VFA dynamic behavior between AT treatment simulated by the model was consistent with the *in vitro* experiments.

### 3.2. Shapley effects - General contribution of the input parameters to the variability of methane and volatile fatty acids production

Figure 8, Figure 9, Figure 10 and Figure 11 display the Shapley effects computed over time for the fourth day of simulation (from 72 to 96h) of 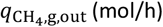, *s*_ac_ (mol/L), *s*_bu_ (mol/L) and *s*_pr_ (mol/L), respectively, for the three dietary scenarios (control, low AT treatment and high AT treatment) studied. This implementation will be able to identify the microbial pathways explaining the variation of CH_4_ and VFA, and to investigate how these pathways change according to the inhibitor doses. Only the IPs associated with a contribution higher than 10% for at least one time step were displayed.

**Figure 8.**
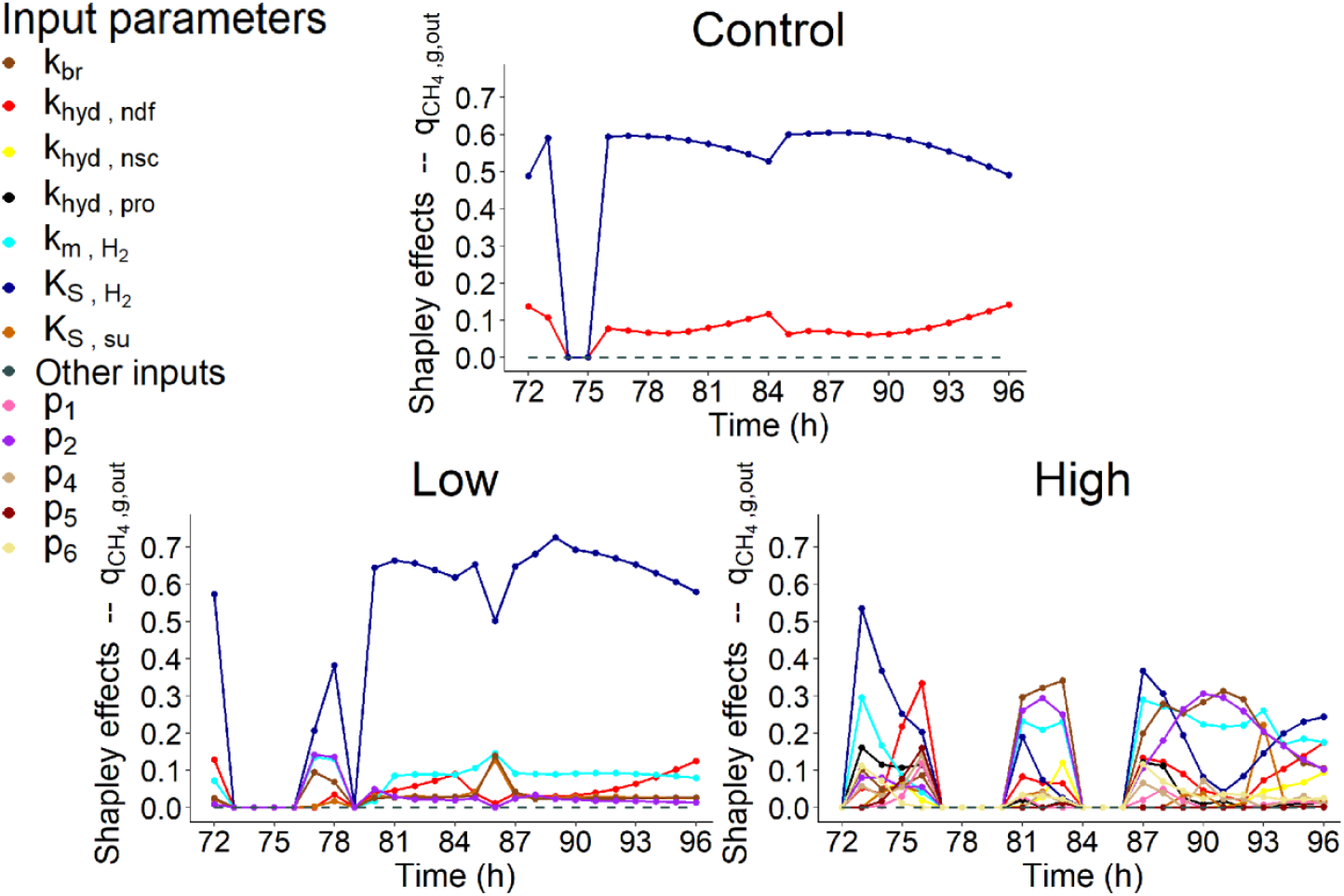
Shapley effects of the influential input parameters (*i*.*e*, parameters with a contribution higher than 10% for at least one time step) over time (h) computed for the fourth day of simulation of the rate of CH_4_ production in gas phase 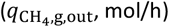 for the three dietary scenarios (control: 0% of *Asparagopsis taxiformis*, low treatment: 0.25% of *Asparagopsis taxiformis* and high treatment: 0.50% of *Asparagopsis taxiformis*).

**Figure 9.**
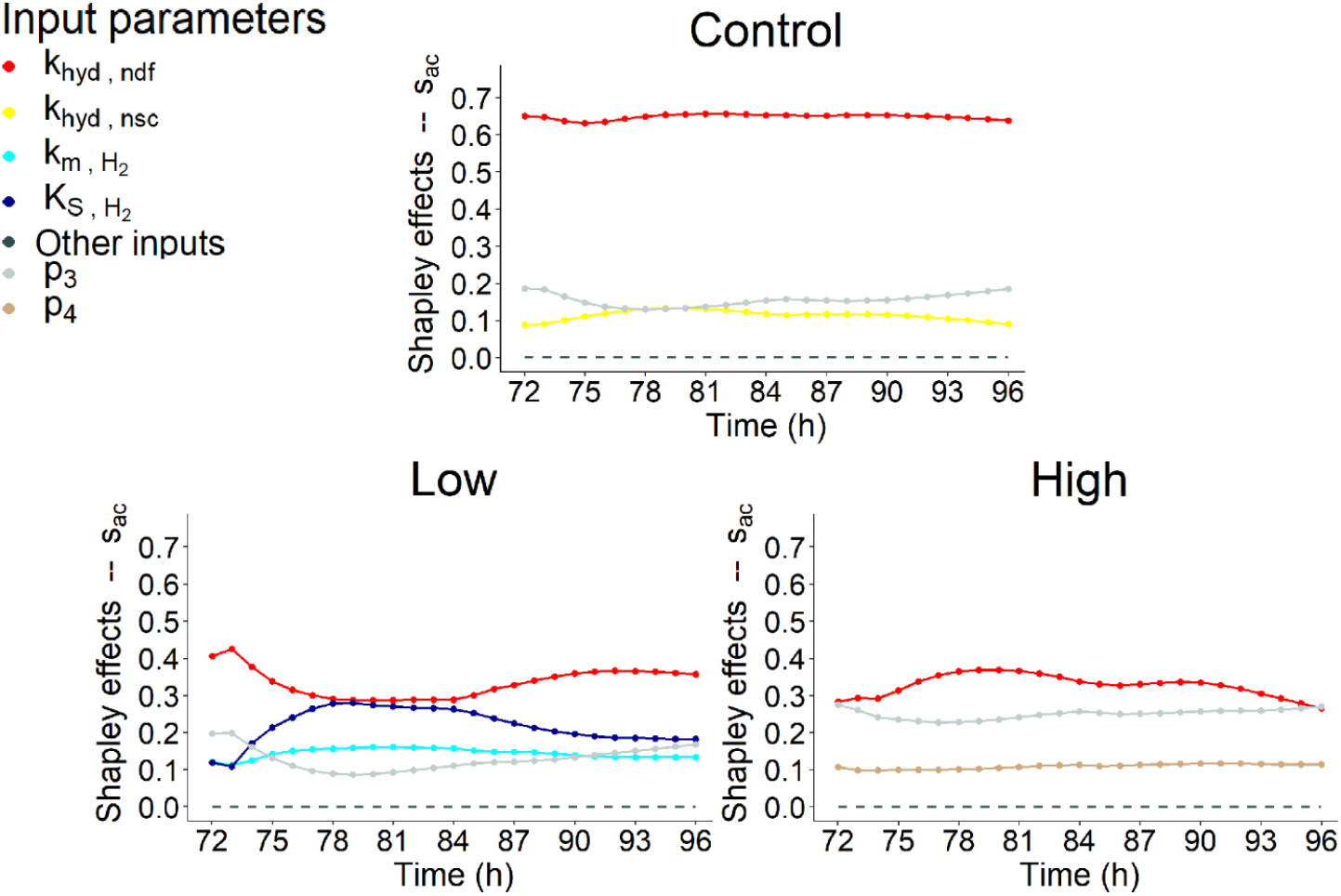
Shapley effects of the influential input parameters (i.e., parameters with a contribution higher than 10% for at least one time step) over time (h) computed for the fourth day of simulation of the acetate concentration (*s*_ac_, mol/L) for the three dietary scenarios (control: 0% of *Asparagopsis taxiformis*, low treatment: 0.25% of *Asparagopsis taxiformis* and high treatment: 0.50% of *Asparagopsis taxiformis*).

**Figure 10.**
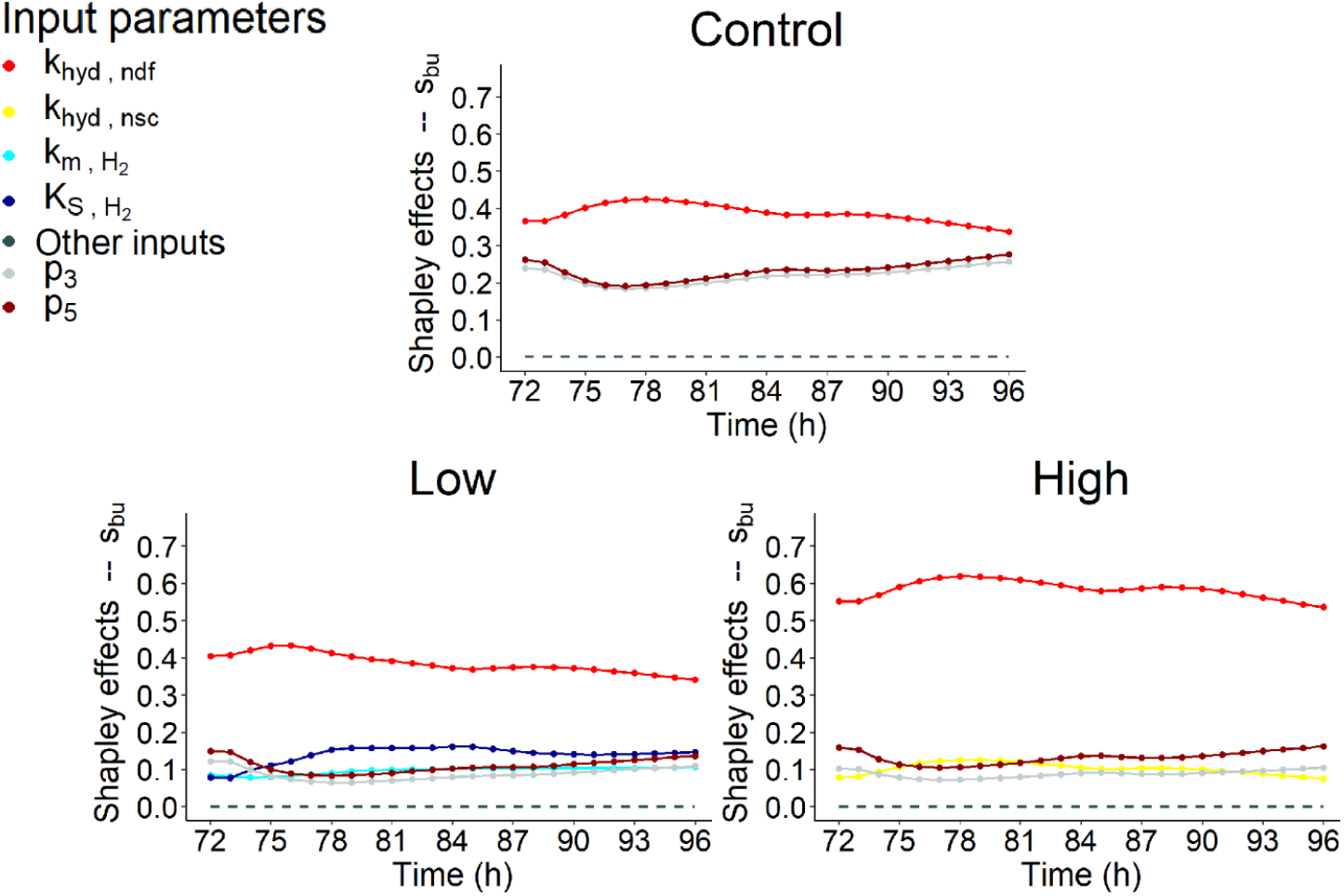
Shapley effects of the influential input parameters (i.e., parameters with a contribution higher than 10% for at least one time step) over time (h) computed for the fourth day of simulation of the butyrate concentration (*s*_bu_, mol/L) for the three dietary scenarios (control: 0% of *Asparagopsis taxiformis*, low treatment: 0.25% of *Asparagopsis taxiformis* and high treatment: 0.50% of *Asparagopsis taxiformis*).

**Figure 11.**
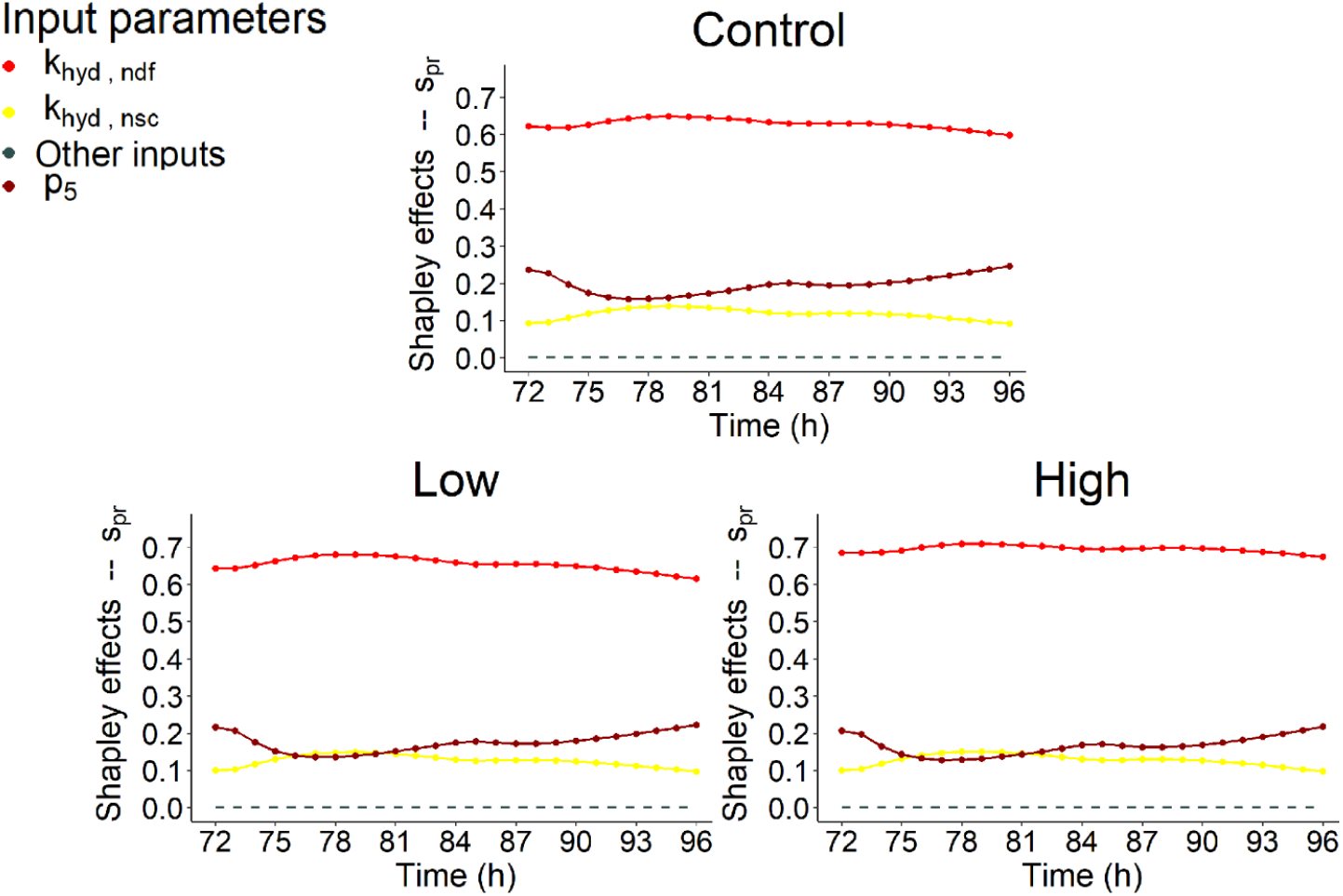
Shapley effects of the influential input parameters (i.e., parameters with a contribution higher than 10% for at least one time step) over time (h) computed for the fourth day of simulation of the propionate concentration (*s*_pr_, mol/L) for the three dietary scenarios (control: 0% of *Asparagopsis taxiformis*, low treatment: 0.25% of *Asparagopsis taxiformis* and high treatment: 0.50% of *Asparagopsis taxiformis*).

For some time steps, the computation led to negative indexes. In this case, the estimates were set to 0. These issues mainly concerned 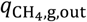 of the two AT treatments and were either due to the outliers in the variability explored in the simulations or to the lack of variability in the simulations for some time steps. The computational time for one dietary scenario was of 24h using the MESO@LR-Platform.

The comparison of our results with those of previous SA conducted on mechanistic models of rumen fermentation is not straightforward given the specific model structures and their mathematical formulation. Consequently, the model structure, and the variables and parameters considered in these models are different from those used in our representation, except for Merk et al. (2023) which conducted its SA on an adapted version of the model of Muñoz-Tamayo et al. (2021). Moreover, regarding the other references than Merk et al., 2023 the comparison of SA results is only valid for the control as other models did not consider AT treatments. Nevertheless, despite these limitations considering our results in relation to those obtained in previous studies provides useful information to improve our knowledge of the whole picture of the rumen fermentation.

#### 3.2.1. Rate of methane production

Figure 8 indicated that the action of the microbial group of hydrogen utilizers represented by the Monod affinity constant 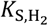, contributed largely the most to the variability of 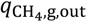 over the fermentation for control and low AT treatment, explaining more than 50% of 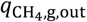 variation over time for both scenarios. The dynamic of the impact of this microbial group was constant over time and slightly followed the dynamic of DMI.

The other influential IP for control was related to microbes degrading the fibers, via the hydrolysis rate constant *k*_hyd,ndf_, highlighting a contribution of c.a. 10% to 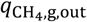 variability with a constant dynamic over time. This IP showed a low influence (c.a. 10%) for low and high AT treatments.

When no dose of AT was considered, Merk et al. (2023) and Huhtanen et al. (2015) also highlighted the impact of fibers degradation component on CH_4_ production variation of Muñoz-Tamayo et al. (2021) and Karoline models, respectively. The initial neutral detergent fiber concentration, which was largely associated with the highest contribution (= 43%) to CH_4_ production variation, and *k*_hyd,ndf_ were the influential IPs in Merk et al. (2023). The other influential IP for the control was related to the flux allocation parameter from glucose utilization to propionate production (*λ*_2_). This last IP was not considered in our SA as we modified the flux allocation parameters in our model version.

For the other SA works carried out under control condition, fat and the degradation of starch and insoluble protein were the other factors associated with an influence on CH_4_ production in Huhtanen et al. (2015). Whereas, van Lingen et al. (2019) highlighted that IPs associated with the fractional passage rate for the solid and fluid fractions in the rumen and NADH oxidation rate explained 86% of CH_4_ predictions variation of a modified version of the Dijkstra model. Our model does not include the different passage rates between solid and liquid fractions. Finally, Dougherty et al. (2017) found that influential IPs on daily CH_4_ production predicted with the AusBeef model were associated with ruminal hydrogen balance and VFA production.

For AT treatments, some IPs associated with the factors modeling the effect of AT on the fermentation were highlighted as influential on 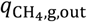 variability. The IPs *p*_2_, which is related to the sigmoid function modeling the direct inhibition effect of AT on methanogenesis, and *k*_br_, which describes the kinetic of bromoform utilization, showed a low (c.a. 10%, at t = from 77 to 78h for *p*_2_, and at t = 87h for *k*_br_) and intermediate (> 20%, at t = from 81 to 83h and from 87 to 93h for both *p*_2_ and *k*_br_) contribution to 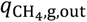 variability for low and high doses of AT, respectively.

Furthermore, AT treatments highlighted the differences of the role of microbial pathways explaining the variation of 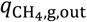 when increasing the dose of AT. When a high dose of AT was supplemented, *k*_br_ and *p*_2_ explained more than 50% of 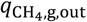 variability in the middle (t = from 81 to 83h) and middle end (t = from 88 to 92h) of the fermentation, replacing a part of the variability explained by 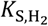. Their influence decreased at the end of the fermentation but was still higher than 20%. When comparing both IPs, *k*_br_ showed a slightly higher influence (< 10%) than *p*_2_. Moreover, other IPs associated with the direct (*p*_1_) or indirect (*p*_4_, *p*_5_ and *p*_6_) effect of bromoform on the fermentations showed a low influence on 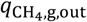 variation for high AT treatment. This highlights that the use of AT to mitigate CH_4_ production led to a shift in the factors associated to microbial pathways of the rumen fermentation impacting the CH_4_ production. The low participation of these IPs to the variability of 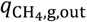 when a low dose of AT was supplemented suggested that the AT dose of this treatment was too low to highlight this shift.

However, the impact of 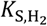 was still important over time for the high AT treatment, especially at the beginning (t = from 73 to 76h) and at the end (t = from 87 to 89h and from 93 to 96h) of the fermentation. Moreover, the hydrogen utilizers microbial group also showed an influence via the maximum specific utilization rate constant 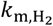 for low and high AT treatments. This influence was low (c.a. 10%) for low AT treatment over the fermentation and was of c.a. 20% or higher at t = 73h, from 81 to 83h and from 87 to 96h for high AT treatment, confirming the importance of this factor on 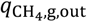 variation.

In presence of a dose of AT (included at a 5% inclusion rate), the high impact of IPs associated with bromoform concentration and the factor *I*_br_ on CH_4_ production variation was also highlighted in Merk et al. (2023). The initial bromoform concentration and *p*_1_ showed the highest contributions (= 46%) to CH_4_ production variation. This study also mentioned the low but non-negligible impact (c.a. 10%) of IPs related to methanogen abundance, total microbial concentration and hydrogen utilizers microbial group, represented by 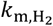. This last IP showed also an influence on 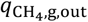 variation for high AT treatment in our work. Therefore, Merk et al. (2023) also identified a shift in the key factors driving CH_4_ production variation in presence of AT.

#### 3.2.2. Volatile fatty acids concentration

Figure 9, Figure 10 and Figure 11 highlighted that similar IPs contributed to the variability of *s*_ac_, *s*_bu_ and *s*_pr_.

VFA concentration variation were highly impacted by fibers degradation represented by the kinetic *k*_hyd,ndf_ for the control and two AT treatments. *k*_hyd,ndf_ was always associated with the highest contribution for *s*_ac_, *s*_bu_ and *s*_pr_ and the three dietary scenarios studied. This result was expected as fiber hydrolysis is the limiting step of the fermentation and VFA proportion. This contribution was very high (> 50%) for *s*_ac_ control, *s*_bu_ high AT treatment and *s*_pr_. While, it was intermediate (between 30 and 40% over the fermentation) for *s*_ac_ low and high AT treatments, and *s*_bu_ control and low AT treatment. The dynamic of this influence was globally constant over time slightly following the dynamic of DMI, except for *s*_ac_ low AT treatment which showed a decrease during the first feed distribution.

Regarding the impact of increasing AT doses on *k*_hyd,ndf_ contribution, it decreased for *s*_ac_ while it was still the most influential IP, with an influence over time varying from 29 to 42% for low AT treatment and from 26 to 37% for high AT treatment.Whereas, it increased for *s*_bu_ and *s*_pr_ with an influence over time, varying from 34 to 42% and 60 to 65% for control, from 34 to 43% and 61 to 68% for low AT treatment and from 54 to 62% and 67 to 71% for high AT treatment, respectively.

Regarding the other influential IPs, hydrogen utilizers microbial group slightly impacted (< 30%) the variation of *s*_ac_ and *s*_bu_ for low AT treatment, with the two IPs representing this group (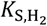 and 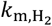). The influence of these IPs was increasing at the beginning of the fermentation, constant in the middle of the fermentation and decreasing at the end of the fermentation. For both variables, this influence was mainly due to 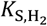, being the second most influential IP from t = 74h to the end of the fermentation with a contribution varying from 17 to 27% over this period of time for *s*_ac_, and explaining a maximum of 16% of the variation of *s*_bu_ (second most influential IP). 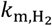 was associated with a low contribution, varying from 11 to 16% over the fermentation for *s*_ac_, and showing a constant dynamic at c.a. 10% for *s*_bu_. Moreover, the degradation of non-fiber compounds with the IP *k*_hyd,nsc_ showed a non-negligible contribution for the control with an influence varying from 13 to 19% for *s*_ac_, for the high AT treatment with an influence of c.a. 10% for *s*_bu_, and for the three dietary scenarios with an influence of c.a. 10% for *s*_pr_.

Furthermore, IPs associated with the functions describing the indirect effect of bromoform on rumen fermentation and quantifying reactions driving flux allocation from glucose utilization to acetate (λ_1_ associated with *p*_3_ and *p*_4_) and propionate (λ_2_ associated with *p*_5_ and *p*_6_) production showed a low impact on VFA concentration variation under our conditions. IP *p*_3_ was associated with an impact on *s*_ac_ and *s*_bu_ variation. This impact was highlighted for the control and two AT treatments studied, with a contribution lower than 20% for control and low AT treatment, which decreased at the beginning of the fermentation and then increased over time, and of c.a. 25% for high AT treatment for *s*_ac_. Whereas, the contribution over time was more important for the control (c.a. 25%) than for the two AT treatments (<20%) for *s*_bu_ with the same dynamic. *p*_4_, the negative slope of λ_1_, slightly impacted *s*_ac_ variation of high AT treatment with a constant dynamic. Regarding λ_2_, *p*_5_ slightly impacted (<30%) *s*_bu_ and *s*_pr_ variation over the fermentation for the three dietary scenarios with a dynamic slightly decreasing at the beginning of the fermentation and then increasing over time, similarly to *p*_3_. The positive slope *p*_6_ did not contribute to VFA concentration variation.

Nevertheless, no shift of the factors associated to microbial pathways impacting VFA production was highlighted when increasing the dose of AT. Moreover, IPs related to the direct effect of bromoform on the fermentation (*p*_1_ and *p*_2_) did not contribute to VFA concentration variation. This suggests that under the conditions evaluated AT had no impact on the biological mechanisms responsible for VFA production, in contrast with the one responsible for CH_4_ production. However, AT supply does have an indirect effect on VFA production to its effect on the lambdas and a variation was observed when considering the molar proportions of VFA (Figure 7). Moreover, new influential IPs were highlighted from a dietary scenario to another for all the VFA, except *s*_pr_.

Morales et al., (2021) studied the sensitivity of 19 IPs on VFA concentration predicted with Molly. It found that the intercept used for rumen pH prediction was the only influential IP, explaining more than 79% of the variation of acetate, butyrate and propionate concentration predictions of Molly. In this study this result was expected as Molly is a whole animal model, which was not the case of our model. No IPs related to rumen pH were considered in our SA, explaining that the selection of this component was not possible in our case.

### 3.3. Uncertainty analysis

The uncertainty of 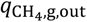 and VFA concentration simulations used to compute the Shapley effects was assessed by studying the variability over time of these simulations. The results were displayed only for 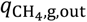.

#### 3.3.1. Rate of methane production

Figure 12 displays the median and quantiles 10% and 90% over time of 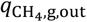 simulations computed by exploring the variability of six factors associated to microbial pathways of the rumen fermentation (fiber carbohydrates, non-fiber carbohydrates, proteins, glucose utilizers, amino acids utilizers, hydrogen utilizers) and four factors of the effect of bromoform on the fermentation (*s*_br_, *I*_br_, *λ*_1_ and *λ*_2_) for the three dietary scenarios.

**Figure 12.**
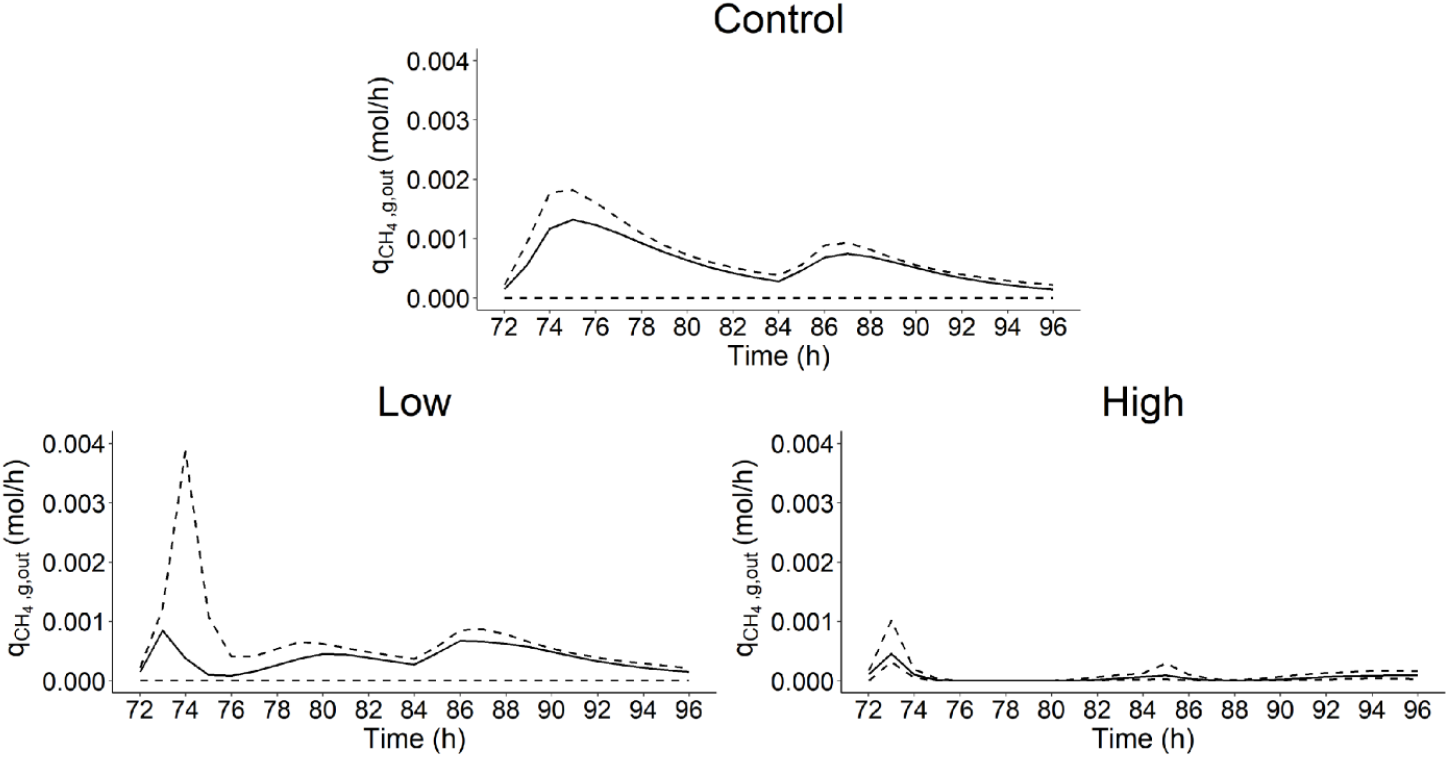
Median and 0.1 and 0.9 quantiles of the rate of methane production in gas phase 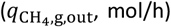 over time (h) computed from the simulations used to calculate the Shapley effects for the three dietary scenarios (control: 0% of *Asparagopsis taxiformis*, low treatment: 0.25% of *Asparagopsis taxiformis* and high treatment: 0.50% of *Asparagopsis taxiformis*).

When not considering the 10% more extreme simulations, the CV highlighted that 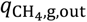 simulations showed a lower variability for low and high AT treatments with a median CV of 0.74 and 0.59, respectively. Whereas, the control showed a median CV of 0.78.

SA results indicated that the variability of 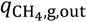 simulations was only explained by the variation of hydrogen utilizers microbial group via 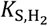 for control and low AT treatment. Whereas, the variability of 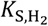 explained an important part, but also with the variability of other factors, of 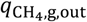 simulation variability for high AT treatment. This suggests that reducing uncertainty associated with 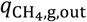 predictions involves to reduce the uncertainty of IPs describing the activity of the hydrogen utilizers microbial group. A way to achieve that is to increase the information available for estimating the variability of parameters describing this microbial group, involving an improvement of our knowledge of it.

Finally, when comparing control and AT treatments, 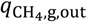 simulation variability was more important for the control than for low and high AT treatments. This indicates that the shift and increase of factors explaining the variation of 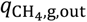 did not lead to an increase of simulation variability, especially for high AT treatment. Merk et al., (2023) found a different result, computing a CV of 0.23 against 1.22 for simulations associated with control and AT treatment, respectively.

However, the range of variation of IPs explored in our study led to outlier simulation for AT treatments. For instance, an IP simulation scenario led to 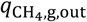 values of 96 and 0.07 mol/h at t = 74 and 73h for low and high AT treatments, respectively. These outliers were not identified for the control. This suggests that some of the range of variation explored for 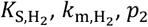 and *k*_br_ was not appropriate when considering AT treatments.

Moreover, Figure 13 highlighted that 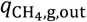 variability varied over time and that this variability was related to the dynamic of DMI (g/h).

**Figure 13.**
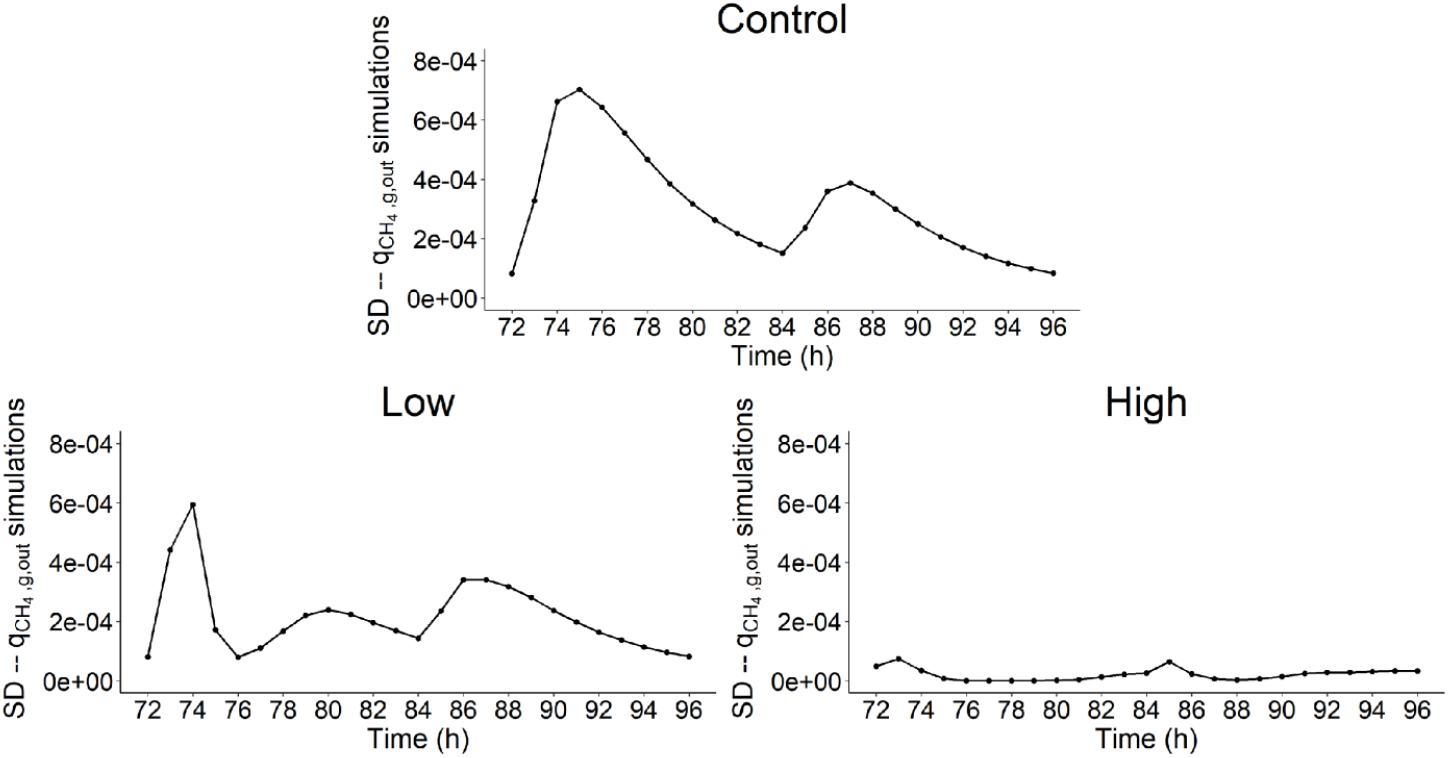
Standard deviation of the simulations of rate of methane production in gas phase 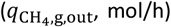 over time (h) used to calculate the Shapley effects for the three dietary scenarios (control: 0% of *Asparagopsis taxiformis*, low treatment: 0.25% of *Asparagopsis taxiformis* and high treatment: 0.50% of *Asparagopsis taxiformis*).

The time periods associated with the highest intake activity (represented by the level of decay of the curve of Figure 2) were between 72 and 78h for the first feed distribution and 84 and 90h for the second feed distribution. The first feed distribution time period was systematically associated with the highest variability of 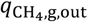 simulations with a maximum SD of 7e-04 mol/h at t = 75h, 6e-04 mol/h at t = 74h and 7e-05 mol/h at t = 73h for control, and low and high AT treatments, respectively. This feed distribution represented 70% of the total DM. The second feed distribution, representing 30% of the total DM, was also associated with an important variability of 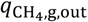 simulations for the three dietary scenarios. Therefore, simulation variability increased during the high intake activity periods of both feed distributions and decreased at the end of it. These results go in line with model developments predicting CH_4_ with dynamic data DMI as single predictor (Muñoz-Tamayo et al., 2019, 2022).

#### 3.3.2. Volatile fatty acids concentration

VFA concentrations were associated with much less variability than 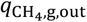. simulations, with a maximum CV of 0.08, 0.12 and 0.09 for *s*_ac_, *s*_bu_ and *s*_pr_, respectively. This suggests that these variables were less sensitive to the variation of factors associated to microbial pathways involved in the rumen fermentation analyzed here. Perhaps the consideration of other parameters such as the yield factors would lead to a higher variability of VFA simulations. The variability of these variables was only explained by the individual variability of the kinetic of fibers degradation *k*_hyd,ndf_. This suggests that the uncertainty related to *k*_hyd,ndf_ measurements generates a low uncertainty on VFA concentrations.

Similarly to 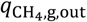, the dynamic of the variability of VFA concentration simulations was related to the dynamic of DMI. This variability increased during the high intake activity periods of the first and second feed distributions, with the highest variability reached for the first feed distribution, and decreased at the end of both feed distributions.

### 3.4. Limitations and perspectives of methods

#### 3.4.1. Sensitivity analysis approaches

The computation of the Shapley effects allowed to identify the influential and non-influential IPs on the variation of 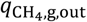 and VFA concentration. This analysis was conducted in a context where the IPs studied were related to factors associated to microbial pathways of the rumen fermentation. Therefore, the aim of the dynamic SA implemented was to gain knowledge on the biological impact of these factors on CH_4_ and VFA production. The use of sensitivity indices for this purpose is becoming increasingly widespread in animal nutrition modeling and our work contributes to this movement. For instance, Merk et al. (2023) conducted local and global SA for identifying key drivers of CH_4_ production with or without AT.

The main originality of our work is the computation of SI over time, leading to a dynamic interpretation of the impact of key drivers on CH_4_ and VFA variation. Moreover, this interpretation was conducted by simulating a typical forage-based diet associated with several realistic dose levels of a highly promising CH_4_ inhibitor. Investigating the dynamic impact of microbial pathways on fermentation outputs, considering diets and treatments applied in the field, is of great interest for animal nutrition, in a context where the combining effect of CH_4_ inhibitors is relevant to study (Muñoz et al., 2024). Integrating the dynamic characteristic is important, as no additive effect of inhibitors was showed under static condition (Muñoz et al., 2024). This highlights that the implementation of dynamic SA on mechanistic models representing the effect of these inhibitors could be used to identify optimal times, frequencies and doses for inhibitor delivery, in order to help develop mitigating strategies using two or more inhibitors. In our work it has not been possible to go that far, but we developed a dynamic SA approach that can be used to reach this aim.

A second originality is the proposition of an approach allowing to discriminate the nature of the contribution of IPs to output variable variation. Although, considering the contribution due to the dependence/correlation between IPs was not relevant in our case study, this work proposes a first methodology to handle this kind of contribution in the case of development of more complex models involving dependent or correlated IPs.

Our study used simulation conditions based on RUSITEC *in vitro* experiments. Future work can use our SA framework to identify useful sampling times and experimental conditions to provide informative data for model refinement in the context of optimal experiment design for parameter estimation.

Regarding our SA results, it is important to mention that they are inherently linked to the representation of the rumen fermentation considered in our case study. Another limitation concerns the range of variation set on IPs studied. Due to the lack of data available, a non-informative distribution was set to explore their variability. By having more information about the variability of these IPs, it will be possible to have more robust results.

#### 3.4.2. Uncertainty analysis

The *in silico* framework used for the SA shows that the factors associated to microbial pathways modeled in our case study mainly impacted CH_4_ prediction uncertainty. This suggests that an improvement in the range of variation of parameters associated with the methanogenesis should lead to a reduction of the uncertainty associated with model predictions. The high AT treatment also showed that the parameters associated with the bromoform effect on the fermentation impacted negatively the prediction uncertainty. These suggestions should be carefully interpreted because limited by the low information available on the numerical values of parameters of the equations representing the rumen fermentation.

## 4. Conclusions

A dynamic sensitivity analysis of a model describing the effect of bromoform (via *Asparagopsis taxiformis*) on rumen fermentation under *in vitro* continuous condition was conducted. The hydrogen utilizers microbial group was identified as the key factor explaining CH_4_ variation over time for the control and low dose treatments. This factor was associated with the microbial methanogenesis. The high AT dose treatment showed a shift in the factors associated to microbial pathways explaining CH_4_ variation, highlighting the emergence of parameters associated with bromoform concentration and direct effect of bromoform on methanogenesis. Moreover, the individual variability of kinetic of fibers degradation explained most of the VFA variation. The uncertainty analysis of simulations computed for SA suggested that reducing the uncertainty of the parameters associated to the kinetics of hydrogen utilizers microbial group should lead to a reduction of model prediction uncertainty. Our work showed that implementing dynamic sensitivity analysis is a promising approach to improve our understanding of mechanisms involved in the rumen fermentation and can help to design optimal experiments assessing CH_4_ mitigation strategies.

## Supporting information

Sobol indices implementation

## 5. Declarations

### Availability of data and material

The scripts for the model simulation and SA are available at Blondiaux (2024).

### CRediT Author contributions

Paul Blondiaux: Conceptualization, Data curation, Formal Analysis, Investigation, Methodology, Software, Writing – original draft

Tristan Senga Kiessé: Conceptualization, Funding acquisition, Methodology, Supervision, Writing – review & editing

Maguy Eugène: Conceptualization, Funding acquisition, Methodology, Supervision, Writing – review & editing

Rafael Muñoz-Tamayo: Conceptualization, Funding acquisition, Investigation, Methodology, Project administration, Software, Supervision, Writing – review & editing

