## Supplementary material for "Dynamic sensitivity analysis of a mathematical model describing the effect of the macroalgae *Asparagopsis taxiformis* on rumen fermentation and methane production under *in vitro* continuous conditions": Sobol indices implementation

This document contain the implementation of the full and independent Sobol indices (Mara et al., 2015) on a mechanistic model of rumen fermentation under *in vitro* continuous conditions accounting for the effect of AT on the fermentation and $CH_4$ production (Muñoz-Tamayo et al., 2021).

### 1. Method

#### 1.1. Definition

Sobol indices (Sobol, 1993) are commonly used when performing SA. The first-order Sobol indice ($S_i$) of an input parameter (IP) $X_i$ quantifies the individual contribution of $X_i$ to output variability, based on the variance decomposition such as

$$S_i = \frac{Var(E[Y|X_i])}{Var(Y)} \quad (1)$$

where Y is the output variable and $X_i$ is an IP. In addition, the total Sobol indice ($T_i$) (Homma and Saltelli, 1996) quantifies the contribution due to the interactions of $X_i$ with other IPs such as

$$T_i = S_i + \sum_{j \neq i} S_{ij} + \sum_{j \neq i, k \neq i, j < k} S_{ijk} + \cdots \quad (2)$$

where $S_i$ is the first-order Sobol indice of $X_i$, $S_{ij}$ is the second-order Sobol indice of $X_i$ quantifying the interactions between $X_i$ and other IPs 2 by 2, $S_{ijk}$ is the third-order Sobol indice of $X_i$ quantifying the interactions between $X_i$ and other IPs 3 by 3.

When dependence/correlation is present among the IPs, the variance decomposition is no longer applicable. Then, $S_i$ and $T_i$ are no longer computable. In this case, Mara et al. (2015) proposed Sobol indices allowing to quantify the contribution of an IP due to its dependence/correlation with other IPs by computing 4 SI:

- the full first-order and total Sobol indices ($S_i^{full}$, $T_i^{full}$) of an IP $X_i$ integrate the dependence/correlation effects between $X_i$ and other IPs;

- the independent first-order and total Sobol indices ($S_i^{ind}$, $T_i^{ind}$) of an IP $X_i$ represent the effects of $X_i$ that are not due to its dependence/correlation with other IPs.

#### 1.2. Interpretation

Sobol indices of Mara et al. (2015) are interpreted in 2 steps:

a. $T_i^{full}$ and $T_i^{ind}$ are compared for quantifying the contribution of $X_i$ due to its dependence/correlation effects.
    - The combination of a high $T_i^{full}$ and a low $T_i^{ind}$ indicates that $X_i$ contributes to output variability only via its dependence/correlation with other IPs;
    - a high $T_i^{ind}$ indicates that $X_i$ contributes to output variability via its own variability and/or its interactions with other IPs;
    - a low $T_i^{full}$ and $T_i^{ind}$ indicates that $X_i$ has no contribution on output variability

b. The second step of the interpretation is the comparison of $S_i^{full}$ and $T_i^{full}$ or $S_i^{ind}$ and $T_i^{ind}$, depending on whether $X_i$ contributes to output variability via its dependence/correlation with other IPs, for quantifying the contribution of $X_i$ due to its interaction effects:
    - $T_i - S_i \approx 0$ indicates that $X_i$ contributes to output variability via its own variability,
    - $T_i - S_i \gg 0$ indicates that $X_i$ contributes to output variability via its interactions with other IPs.

#### 1.3. Numerical computation

For estimating the 4 indices, IPs were sampled using the Sobol sequences, which were identified as providing a better stability for Sobol indices estimation (Blondiaux et al., 2022). 3000 simulations and 100 bootstrap replications using the Sobol (Sobol, 1993) and Saltelli (Saltelli et al., 2008) estimators for computing first-order ($S_i^{full}$ and $S_i^{ind}$) and total ($T_i^{full}$ and $T_i^{ind}$) indices, respectively, were conducted with the R package "sensobol" (Puy et al., 2022), leading to 54000 model evaluations. Moreover, the estimation of conditional densities of the IPs is required for modelling their dependence structure. This estimation was conducted using a vine copula model.

In addition to the Shapley effects, the interpretation of $S_i^{full}$, $T_i^{full}$, $S_i^{ind}$ and $T_i^{ind}$ allows a distinction of the individual, interaction and dependence/correlation effects. Therefore, in our work, both methods were used in a complementary approach. First, the Shapley effects of the 16 IPs studied were computed for estimating their percentage of contribution, integrating the 3 effects, to the variability of output variables of interest. Second, the full and independent Sobol indices were computed for identifying what is the main source of contribution among these 3 effects.

### 2. Results and discussion

Similarly to the Shapley effects, the full and independent Sobol indices of the 16 IPs were computed over time, considering the 4th day of simulation, for the 4 output variables and 3 dietary scenarios studied. These indices were computed in addition to the Shapley effects for identifying the nature of the contribution of the IPs to output variable variation. They allowed us to investigate if the influential IPs contributed through their own variability or through the variability generated by their dependence/correlation and/or their interactions with other IPs. The computational time for one dietary scenario was of 5h using the MESO@LR-Platform.
This step was done as an academic exercise since, by construction, the model does not have dependency between IPs. This means that the difference between $\mathrm{T_i^{full}}$ and $\mathrm{T_i^{ind}}$ was systematically close to 0 for all the IPs, output variables and dietary scenarios studied, indicating that the contribution of IPs to output variable variation were not due to the dependence or correlation between IPs. However, the methodology here illustrated can be useful to address the dependency aspects in future model extensions.

#### 2.1. Rate of methane production

The full and independent Sobol indices were displayed only for $q_{\mathrm{CH_4,g,out}}$ (mol/h) (Figure 1).

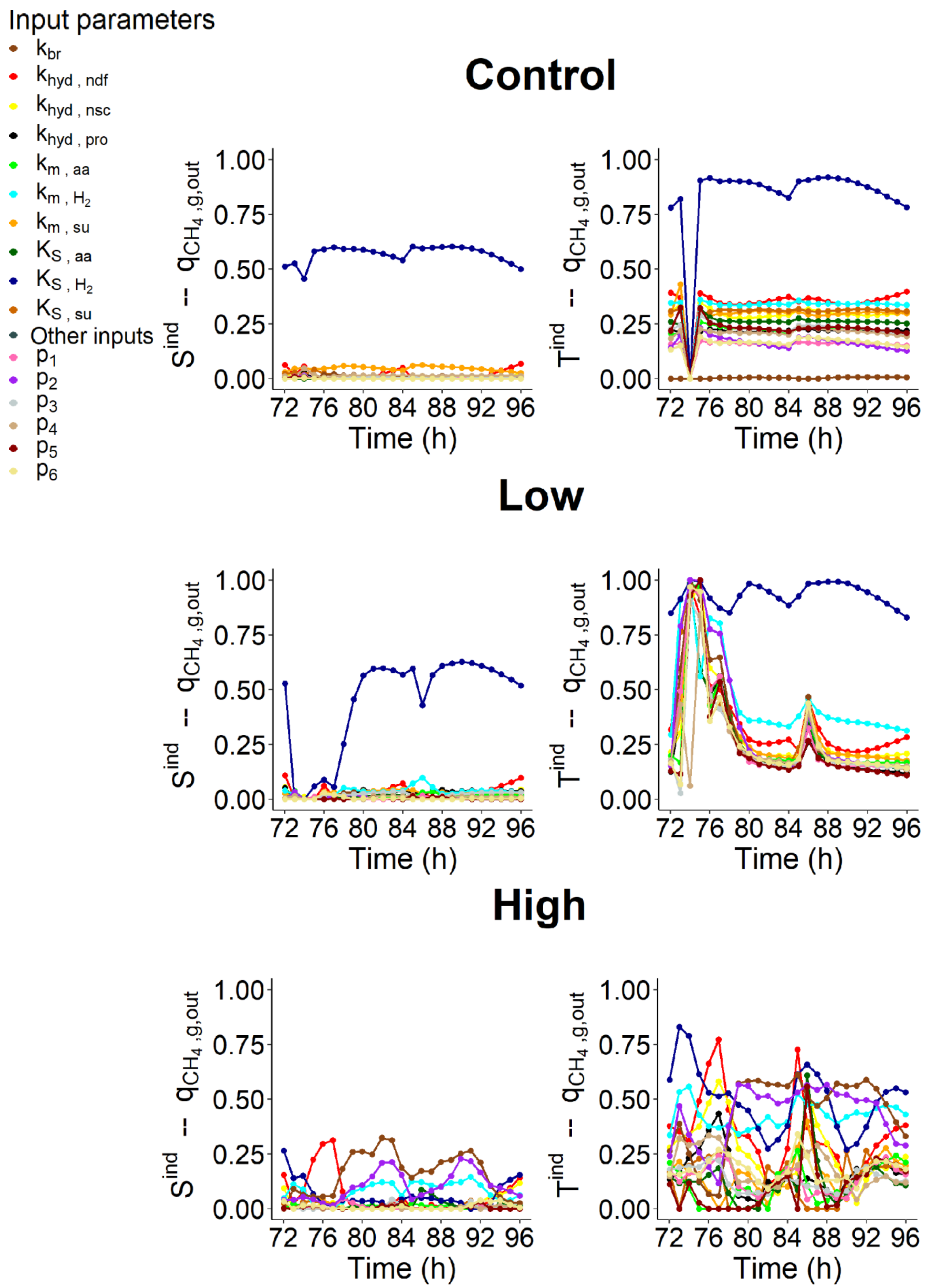


**Figure 1**. Independent first-order ($S^{ind}$) and total Sobol indices ($T^{ind}$) of the input parameters over time (h) computed for the 4th day of simulation of the rate of $CH_4$ production in gas phase ($q_{CH_4,g,out}$, mol/h) for the 3 dietary scenarios (control: 0% of *Asparagopsis taxiformis*, low treatment: 0.25% of *Asparagopsis taxiformis* and high treatment: 0.50% of *Asparagopsis taxiformis*).

89

The difference between $\mathrm{T_i^{ind}}$ and $\mathrm{S_i^{ind}}$ over time indicated that interactions between IPs had an important influence on its variation for the control and two AT treatments over the fermentation.

For control, the variability of $K_{\mathrm{S,H_2}}$ alone had a high contribution on the variation of $q_{\mathrm{CH_4,g,out}}$ ($\mathrm{S}_{K_{\mathrm{S,H_2}}}^{\mathrm{ind}}$ varying from 45 to 60%). The contribution of $K_{\mathrm{S,H_2}}$ increased over the fermentation when considering its influence via the interactions with other IPs over time ($\mathrm{T}_{K_{\mathrm{S,H_2}}}^{\mathrm{ind}}$ varying from 78 to 92%). The other IPs did not show any contribution due to their own variability ($\mathrm{S}_{i,\mathrm{i}\neq K_{\mathrm{S,H_2}}}^{\mathrm{ind}}$ c.a. 0). However, their independent total indices indicated that all the IPs interacted with $K_{\mathrm{S,H_2}}$ ($\mathrm{T_i^{ind}} \geq 10\%$), except $k_{\mathrm{br}}$ with $\mathrm{T}_{k_{\mathrm{br}}}^{\mathrm{ind}}$ c.a. 0 over the fermentation (which is obvious since there is not bromoform in the control). The influential contribution due to the interactions between IPs was mostly due to interactions of $K_{\mathrm{S,H_2}}$ with other IPs such as the fibers degradation ($k_{\mathrm{hyd,ndf}}$), the maximum utilization rate of hydrogen utilizers microbial group ($k_{\mathrm{m,H_2}}$), the microbial group of sugars utilizers ($K_{\mathrm{S,su}}$ and $k_{\mathrm{m,su}}$), the non-fibers degradation ($k_{\mathrm{hyd,nsc}}$) and the microbial group of amino acids utilizers ($K_{\mathrm{S,aa}}$) with on average $\mathrm{T}_{\mathrm{i,i}=k_{\mathrm{hyd,ndf}},k_{\mathrm{m,H_2}},K_{\mathrm{S,su}},k_{\mathrm{m,su}},k_{\mathrm{hyd,nsc}},K_{\mathrm{S,aa}}}^{\mathrm{ind}}$ higher than 25% over time.

For low AT treatment, $K_{\mathrm{S,H_2}}$ was the only IP contributing to $q_{\mathrm{CH_4,g,out}}$ variation via its own variability over time, similarly to the control. $S_{K_{\mathrm{S,H_2}}}^{\mathrm{ind}}$ varied from 0 to 63% over the fermentation with a contribution lower than 10% from t = 73 to 77h, corresponding to the highest intake activity of the first feed distribution. During this time period, $q_{\mathrm{CH_4,g,out}}$ variation was only explained by the interactions between the 16 IPs, with $\mathrm{T_i^{ind}}$ higher than 80% for all the IPs for at least one time step during this time period. This highlights the importance of quantifying the interactions between IPs in SA approaches used. Similarly to the control, $K_{\mathrm{S,H_2}}$ also explained almost all the variability of $q_{\mathrm{CH_4,g,out}}$ over the fermentation when considering the contribution via the interactions with other IPs with $\mathrm{T}_{K_{\mathrm{S,H_2}}}^{\mathrm{ind}}$ varying from 82 to 100% over time. These interactions mainly included $k_{\mathrm{m,H_2}}$ and $k_{\mathrm{hyd,ndf}}$ with $\mathrm{T}_{k_{\mathrm{m,H_2}}}^{\mathrm{ind}} = 36\%$ and $\mathrm{T}_{k_{\mathrm{hyd,ndf}}}^{\mathrm{ind}} = 28\%$ on average over time.

For high AT treatment, $\mathrm{S_i^{ind}}$ was lower than 32% for all the IPs, indicating that the variation of $q_{\mathrm{CH_4,g,out}}$ was mainly due to the interactions between IPs. The IP with the most important individual contribution over the fermentation was $k_{\mathrm{hyd,ndf}}$ during the first feed distribution ($\mathrm{S}_{k_{\mathrm{hyd,ndf}}}^{\mathrm{ind}}$ varying from 22 to 31% from t = 75 to 77h) and at the end of the fermentation ($\mathrm{S}_{k_{\mathrm{hyd,ndf}}}^{\mathrm{ind}} = 14\%$ at t = 96h). Then, the contribution of other IPs such as $k_{\mathrm{br}}$ ($\mathrm{S}_{k_{\mathrm{br}}}^{\mathrm{ind}}$ varying from 14 to 32%), $p_2$ ($\mathrm{S}_{p_2}^{\mathrm{ind}}$ varying from 4 to 23%) and $k_{\mathrm{m,H_2}}$ ($\mathrm{S}_{k_{\mathrm{m,H_2}}}^{\mathrm{ind}}$ varying from 3 to 14%) was low in the middle of fermentation from t = 78 to 92h. Also, $K_{\mathrm{S,H_2}}$ showed the most important contribution at the beginning and end of the fermentation with $\mathrm{S}_{K_{\mathrm{S,H_2}}}^{\mathrm{ind}} = 26$ and $15\%$ at t = 72 and 96h, respectively. When considering the interactions between IPs, their contribution increased significantly. All the IPs showed a non-negligible contribution (> 20%) for at least one time step, indicating that the 16 IPs were considered in the interactions contributing to

$q_{\mathrm{CH_4,g,out}}$ variation. IP $K_{\mathrm{S,H_2}}$ contributed the most via its interactions with an average difference between $\mathrm{T}^{\mathrm{ind}}_{K_{\mathrm{S,H_2}}}$ and $\mathrm{S}^{\mathrm{ind}}_{K_{\mathrm{S,H_2}}}$ over time of 42%. IP $k_{\mathrm{m,H_2}}$ also showed an important contribution throughout the fermentation via the interactions with other IPs with an average constant contribution of 34% over time. The third and fourth IPs contributed the most via the interactions were $p_2$ and $k_{\mathrm{br}}$, respectively, with on average $\mathrm{T}^{\mathrm{ind}}_{p_2} - \mathrm{S}^{\mathrm{ind}}_{p_2} = 33\%$ and $\mathrm{T}^{\mathrm{ind}}_{k_{\mathrm{br}}} - \mathrm{S}^{\mathrm{ind}}_{k_{\mathrm{br}}} = 28\%$ over time. These contributions were more important in the middle of fermentation from t = 78 to 92h. Finally, $k_{\mathrm{hyd,ndf}}$ and $k_{\mathrm{hyd,nsc}}$ also showed a non-negligible contribution via the interactions with on average $\mathrm{T}^{\mathrm{ind}}_{k_{\mathrm{hyd,ndf}}} - \mathrm{S}^{\mathrm{ind}}_{k_{\mathrm{hyd,ndf}}} = 28\%$ and $\mathrm{T}^{\mathrm{ind}}_{k_{\mathrm{hyd,nsc}}} - \mathrm{S}^{\mathrm{ind}}_{k_{\mathrm{hyd,nsc}}} = 24\%$ over time.

The results highlighted that the contribution of interactions between IPs was higher for AT treatments than for the control, with the highest contribution for low AT treatment. Merk et al. (2023) also identified the importance of the contribution due to the interactions between IPs for AT treatments. However, no impact of the interactions was highlighted for the control. The interactions between hydrogen utilizers microbial group via $K_{\mathrm{S,H_2}}$ and the other factors associated to microbial pathways of the rumen fermentation considered had an intermediate impact (~25%) on $q_{\mathrm{CH_4,g,out}}$ variation for the 3 dietary scenarios considered. Most of the time, these interactions included all the IPs, except $k_{\mathrm{br}}$ for the control.

The inclusion of all the IPs in the interactions impacting $q_{\mathrm{CH_4,g,out}}$ variation suggests that the model should be improved to better characterize the interactions. The incorporation of microbial genomic knowledge is expected to improve the representation of rumen microbial fermentation in mathematical models (Davoudkhani et al., 2024; Muñoz-Tamayo et al., 2023).

#### 2.2. Volatile fatty acids concentration

For VFA concentrations ($s_{\mathrm{ac}}$, $s_{\mathrm{bu}}$ and $s_{\mathrm{pr}}$, mol/L), the contribution of the interactions between IPs to their variation was lower than 20% (i.e. $\mathrm{T}^{\mathrm{ind}}_{\mathrm{i}} - \mathrm{S}^{\mathrm{ind}}_{\mathrm{i}} \leq 20\%$) for the 3 dietary scenarios studied, except for $p_5$ which described the fraction of glucose utilized to produce propionate and showed a strong influence on $s_{\mathrm{ac}}$, $s_{\mathrm{bu}}$ and $s_{\mathrm{pr}}$ variation of the control with a maximum contribution of 60, 38 and 29%, respectively. For these 3 variables, the maximum contribution of $p_5$ via the interactions was reached at the end of the fermentation. Also, the interactions between $k_{\mathrm{m,H_2}}$ and other IPs showed a contribution of 24% and 19% to $s_{\mathrm{ac}}$ variation for low and high AT treatments, respectively.
